# Cardiac Magnetic Resonance Studies in a Large Animal Model that Simulates the Cardiac Abnormalities of Human Septic Shock

**DOI:** 10.1101/2024.02.05.578971

**Authors:** Verity J. Ford, Willard N. Applefeld, Jeffrey Wang, Junfeng Sun, Steven B. Solomon, Stanislav Sidenko, Jing Feng, Cynthia Sheffield, Harvey G. Klein, Zu-Xi Yu, Parizad Torabi-Parizi, Robert L. Danner, Vandana Sachdev, Michael A. Solomon, Marcus Y. Chen, Charles Natanson

## Abstract

**Background:** Septic shock, in humans and in our well-established animal model, is associated with increases in biventricular end diastolic volume (EDV) and decreases in ejection fraction (EF). These abnormalities occur over 2 days and reverse within 10 days. Septic non-survivors do not develop an increase in EDV. The mechanism for this cardiac dysfunction and EDV differences is unknown.

**Methods:** Purpose-bred beagles randomized to receive intrabronchial *Staphylococcus aureus* (n=27) or saline (n=6) were provided standard ICU care including sedation, mechanical ventilation, and fluid resuscitation to a pulmonary arterial occlusion pressure of over 10mmHg. No catecholamines were administered. Over 96h, cardiac magnetic resonance imaging, echocardiograms, and invasive hemodynamics were serially performed, and laboratory data was collected. Tissue was obtained at 66h from six septic animals.

**Results:** From 0-96h after bacterial challenge, septic animals *vs.* controls had significantly increased left ventricular wall edema (6%) and wall thinning with loss of mass (15%) which was more pronounced at 48h in non-survivors than survivors. On histology, edema was located predominantly in myocytes, the interstitium, and endothelial cells. Edema was associated with significantly worse biventricular function (lower EFs), ventricular-arterial coupling, and circumferential strain. In septic animals, from 0-24h, the EDV decreased from baseline and, despite cardiac filling pressures being similar, decreased significantly more in non-survivors. From 24-48h, all septic animals had increases in biventricular chamber sizes. Survivors biventricular EDVs were significantly greater than baseline and in non-survivors, where biventricular EDVs were not different from baseline. Preload, afterload, or HR differences did not explain these differential serial changes in chamber size.

**Conclusion:** Systolic and diastolic cardiac dysfunction during sepsis is associated with ventricular wall edema. Rather than differences in preload, afterload, or heart rate, structural alterations to the ventricular wall best account for the volume changes associated with outcome during sepsis. In non-survivors, from 0-24h, sepsis induces a more severe diastolic dysfunction, further decreasing chamber size. The loss of left ventricular mass with wall thinning in septic survivors may, in part explain, the EDV increases from 24-48h. However, these changes continued and even accelerated into the recovery phase consistent with a reparative process rather than ongoing injury.

**Clinical Perspective:** *What is new?:*

- Utilizing multimodal imaging and hemodynamics, we demonstrate the cardiac changes of sepsis have injury and reparative phases.
- The injury phase (0-24h) has EDV decreases more profound in non-survivors and is associated with worse ventricular compliance, myocardial edema, and diastolic dysfunction.
- The recovery phase has left ventricular mass loss with wall thinning in survivors that explains the EDV increases (24-96h). These progressed into the EF recovery phase consistent with a reparative process removing damaged tissue.
- This is the first controlled CMR sepsis study supporting ventricular wall edema is a fundamental aspect of sepsis pathophysiology and dry mass loss a reparative mechanism.

*What are the clinical implications?:*

- Despite optimizing filling pressures, the cardiac changes in ventricular wall structure and function associated with survival and non-survival in sepsis still occurred, thereby discounting fluid resuscitation as the major factor of therapeutic importance for cardiac function and survival.
- The changes reported here have potential implications for sepsis treatment especially in the field of fluid resuscitation. These findings yield new understanding into the pathophysiology of sepsis cardiac dysfunction and allow for novel phenotyping and prognosticating of the syndrome with ventricular compliance and EDVs. This also offers potentially high yielding targets for research for new therapeutic approaches for sepsis and heart failure.

## Introduction

Understanding cardiac function in sepsis remains constrained by unexplained findings. Historically, sepsis was thought of as a “high output” state; yet, in a substantial number of patients, cardiac output was found to be depressed.^1–3^ In the 1980s, serial measurements of left ventricular ejection fraction (LVEF) via radionucleotide cine angiocardiograms (RNCA) coupled with data from indwelling thermodilution pulmonary artery catheters (PACs) allowed detailed assessments of myocardial function during septic shock.^4–6^ Human and animal data demonstrated in survivors, a fall in LVEF over two days after the onset of shock (humans) or bacterial challenge (animal models) with recovery to near normal in 7-10 days.^4–6^ Simultaneous use of RNCA and PAC measurements provided a calculated Left Ventricular End Diastolic Volume (LVEDV) equivalent to stroke volume (SV)/LVEF. In survivors, the LV chamber size markedly increased during the first two days after the onset of shock and normalized as the LVEF recovered over 7-10 days.^4–6^ What remains unexplained in humans and animal survivors is the mechanism of this transient, rapidly reversible cardiac dysfunction and why septic human^4, 5^ and animal^7^ non-survivors do not undergo a similar degree of LV dilation.

This pattern, of transient LVEF decreases and LVEDV increases in septic animal survivors which occurs over 7-10 days, remains immutable despite altering the type,^8^ viability,^8^ dose,^7^ site of bacterial inoculation,^6, 9, 10^ administration of intravenous proinflammatory mediator challenges,^11^ or increasing doses of mediator challenge.^12^ This suggests a common pathway of cardiac injury for disparate inflammatory insults to the host. Yet, in these animals at day two when the LVEF falls, and LV dilation is most profound, light, and electronic microscopy (EM) do not show evidence of myocardial inflammatory cell infiltrates. However, there is interstitial, myocyte, and endothelial cell edema consistent with a non-occlusive diffuse micro-vascular injury with fibrin deposition. While mild focal myofilament dropout is observed, gross abnormities in myofilament structure or frank myocyte loss are not seen.^13^ Since the 1980s, various observational transthoracic echocardiogram (TTE) studies in septic humans have mostly confirmed these cardiac LVEF and LVEDV findings.^14–16^ Despite the prognostic and potential therapeutic implications of these myocardial ventricular volume changes, the underlying pathophysiology has been largely unexplained.

We utilized, serial cardiac magnetic resonance (CMR) imaging in a septic animal model to obtain precise direct measurements of both Right Ventricular End Diastolic Volume (RVEDV) and LVEDV at crucial time points. Simultaneously, we obtained invasive hemodynamic data to examine if sepsis-induced changes in preload, afterload, or heart rate contributed to any differential changes found in cardiac chamber size related to survival. To explore the contribution of changes within the cardiac walls to alterations in ventricular volumes, serial CMR measures of edema and LV mass were obtained. Tissue samples for light and electron microscopy (EM) were acquired to look for an etiology of the cardiac dysfunction and to correlate the changes we observed on imaging with morphological changes on pathology.

## Methods

Thirty-three purpose-bred beagles (9-15 kg, 18-30 mo. old, male, Marshall Farms) were studied. On day one (baseline), 27 tracheostomized, sedated and mechanically ventilated animals received an intrabronchial challenge of *S. aureus* (0.5 - 1.0 ×10^9^ Colony Forming Units/kg) while six animals received an equivalent Phosphate Buffer Solution (PBS) challenge. The *S. aureus* was prepared and administrated as previously described.^10^

Animals were monitored around the clock for 96h to simulate care in a medical as well as an animal hospital intensive care unit as previously described.^10^ Ceftriaxone (50mg/kg IV) starting 4h after intrabronchial challenge was administered q24h. There were no catecholamine infusions given. Maintenance fluid (2ml/kg/h of Normasol-M with 5% dextrose supplemented with KCl (27mEq/l)) were infused 0-96h. Additionally, a PAC was placed before intrabronchial challenges, and the pulmonary artery occlusion pressure (PAOP) was measured q4h throughout. Whenever the PAOP fell below 10mmHg, a 20ml/kg bolus of plasmalyte (Vetivex) was given over 20min and repeated until achieving a PAOP >10mmHg. At 62h, six of the septic animals were sacrificed and tissue was obtained. At 96h, animals that did not succumb to sepsis were deemed survivors and euthanized.^10^ Throughout, during mechanical ventilation, animals had their fractional inspired oxygen levels, positive end expiratory pressure levels, tidal volumes and ventilatory rates adjusted to maintain normal arterial oxygen and carbon dioxide levels as per protocol.^10^ Animals throughout also received sedation titrated to physiological endpoints, stress ulcer and venous thromboembolism prophylaxis, and their position changed at set intervals to avoid stasis ulcers.^10^ All animals were treated equally, except for the type of intrabronchial challenge. The study protocol was reviewed and approved by the NIH, Critical Care Institutional Animal Care and User Committee (CCM19-04).

Femoral lines, PACs and a tracheostomy were placed under anesthesia and baseline TTEs, CMR exams and blood work were obtained prior to intrabronchial challenges.^10^ TTEs were done daily, and CMR exams were performed at two and four days after challenges. At multiple timepoints, laboratory parameters were obtained using arterial blood for ABGs, CBCs, and serum chemistries (Heska). Cytokines and chemokines were measured using commercially available ELISA kits (ProcartaPlex, Invitrogen, ThermoFisher, Waltham, MA). Troponin I and BNP was determined using a multidetector microplate reader (Synergy HT, BioTek Instruments, Winooski, VT).

Please see e-supplementary **Methods** for details on TTE, CMR, literature searches, histology, animal inclusion criteria and statistical analysis.

## Results

At 48 and 96h after challenge, septic animals had significant worsening from baseline of mean Right Ventricular Ejection Fraction (RVEF), Right Ventricular-Pulmonary Artery coupling (RV-PA), and circumferential strain (Figure 1, Panel A, B, C, respectively), as well as increases in mean Right Ventricular End Systolic Volume (RVESV) and RVEDV (Panel D&E respectively) and decreases in mean Right Ventricular Stroke Volume (RVSV) (48h only, Panel F). Non-septic controls had no significant changes in these parameters throughout. The changes from baseline in mean RVEF, RVESV, RVSV, and RV-PA coupling of septic animals were significantly more profound than in controls. The corresponding left heart parameters had remarkably similar but somewhat less pronounced abnormalities (Panel G-L) potentially related to the significantly reduced afterload on the left *versus* right-sided circulation. Septic animals *vs.* controls from 0-96h developed significantly higher mean mPAPs and Pulmonary Vascular Resistances (PVR), and lower mean MAPs and Systemic Vascular Resistances (SVR) (e-supplementary Figure 1, Panel A-D). The quantity of fluids 0-96h received and cardiac filling pressures (PAOP) were similar comparing septic animals *vs.* controls (all, p >0.05) (Panel E-H). Consequently, there were no preload differences to explain the cardiac dysfunction observed in septic animals *vs.* controls. Further, the right and left-sided circulations demonstrated opposite hemodynamic trends of afterload which cannot explain the similar biventricular dysfunction in sepsis.

**Figure 1:**
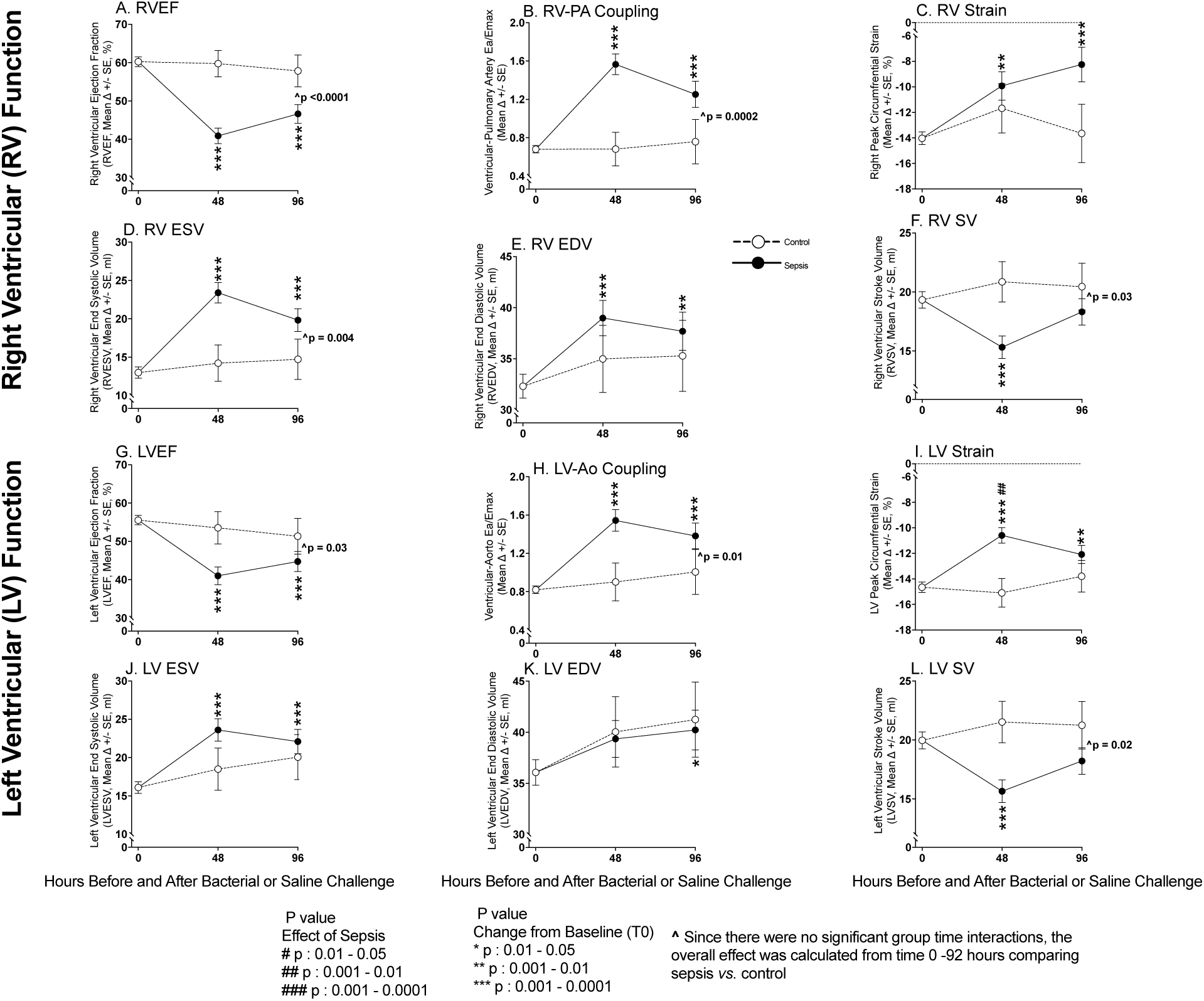
Serial Cardiac MRI Imaging During Experimental Septic Shock. Serial cardiac changes from baseline to 96h plotted from a common origin of the mean values of all animals at time 0 during sepsis (filled circles) *vs.* control (open circles) in the right (Top Panels A-F) and left (Bottom Panel G-L) ventricles as ascertained by CMR.

The importance of these CMR findings to outcome was investigated (Figure 2). The increase 0-48h in mean RVEDV (Panel A) and RVESV (Panel B) was significantly greater in septic survivors *vs.* non-survivors, who had no significant changes from baseline. The mean RVSV (Panel C) in both survivors and non-survivors at 48h similarly significantly decreased. In both survivors and non-survivors at 48h, there was similar significant worsening in mean RVEF (Panel D). Mean RV circumferential strain diminished significantly in survivors at 48h but there was no significant change from baseline in non-survivors (Panel E). In both survivors and non-survivors at 48h, there was similar significant worsening in mean RV-PA coupling (Panel F). Similar findings were observed in the left-sided circulation except for LV strain where there was less of a difference between survivors and non-survivors (Panel G-L). Thus, the only parameter that clearly differentiated septic survivors from non-survivors was changes in biventricular volumes.

**Figure 2:**
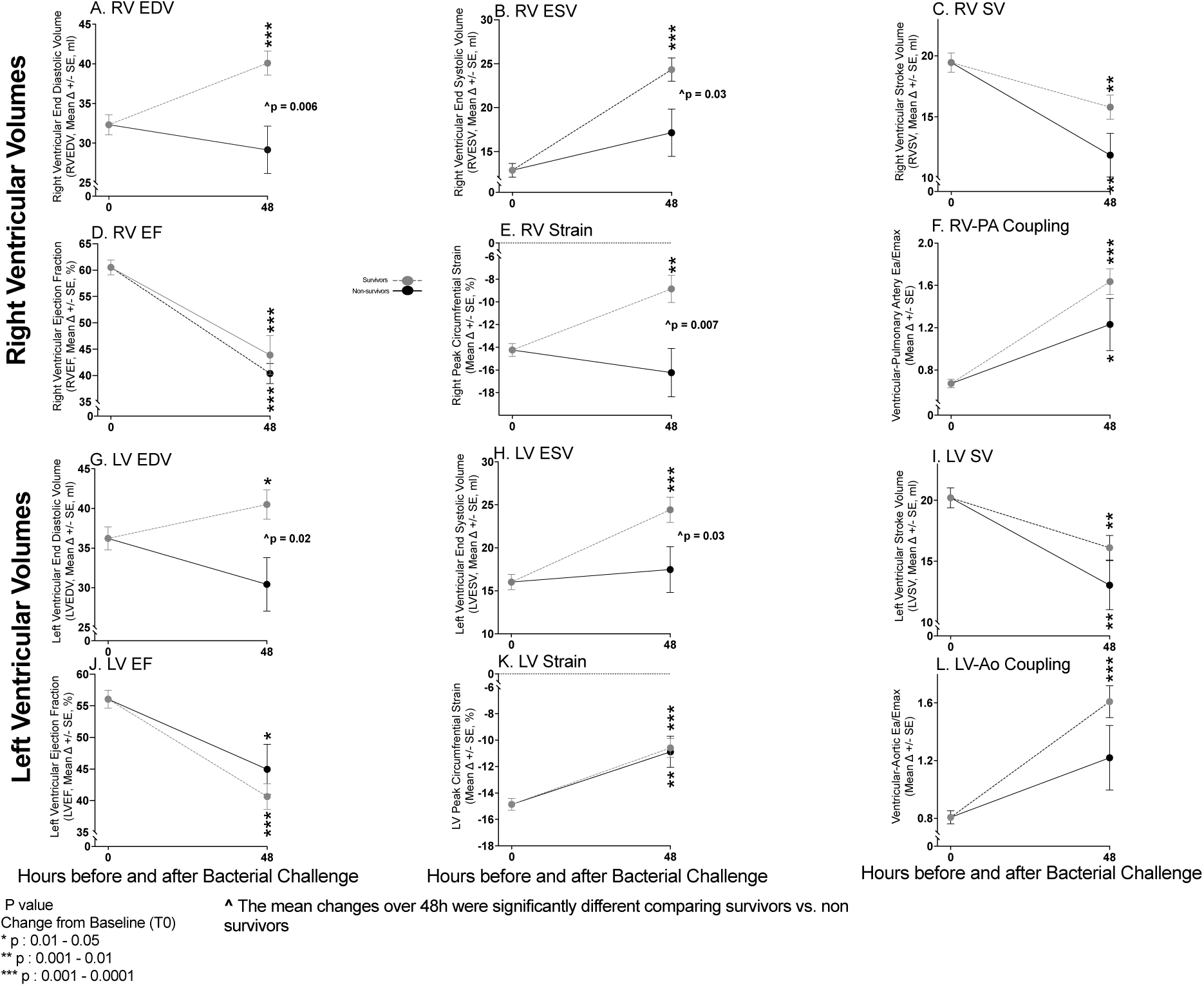
Serial Cardiac MRI Imaging Comparing Septic Survivors and Non-survivors. At 48h, changes from baseline in the heart ascertained by CMR plotted from a common origin of mean values of septic animals at time 0 in septic survivors (grey circles) *vs.* septic non-survivors (filled circles) in the right (Top Panel A-F) and left (Bottom Panel G-L) ventricles.

We next examined if loading conditions could explain the increases in chamber sizes at 48h in septic survivors (Figure 3). There were no significant differences in mean cardiac filling pressures between septic survivors *vs.* non-survivors from 0-48h associated with these differences in volumes either in the left-sided circulation (PAOP) pre- (e-supplementary Figure 2, Panel A) or post completion of the fluid boluses algorithm (Figure 3, Panel A) or the right-sided (Central Venous Pressure, CVP) circulation (Figure 3, Panel B). Ventricular compliance plots showing mean changes over time in ventricular filling pressures *vs.* volumes are shown in Figure 3, Panel C for the LV and in Panel D for the RV. These indicate that marked biventricular compliance differences explain the ventricular volume differences between survivors and non-survivors. A significant shift from baseline to 48h to the right and upward occurred in survivors with increases in both mean PAOP (LV filling pressure) and CVP (RV filling pressure) and concomitant increases in both mean LVEDV (Panel C) and RVEDV (Panel D), respectively. A similar shift to the right with increases in ventricular filling pressures (both mean PAOP and CVP) occurred in non-survivors but no concomitant significant shift upward took place (i.e., no increases in biventricular volumes). Moreover, in both survivors and non-survivors, there was no relationship between changes in filling pressures [PAOP (Figure 3 Panel E) and CVP (Figure 3 Panel F)] from baseline to 48h and changes in LVEDV and RVEDV. Thus, in survivors and non-survivors alike, the differential changes in ventricular volumes were not associated with corresponding alterations in filling pressures. The lack of linearity in the relationship between pressure and volume suggests that changes in ventricular size are not simply the result of fluid administration and sepsis-induced alterations in vascular tone. Rather, in non-survivors, failure to increase ventricular volumes appears intrinsic to the walls themselves. Decreases in wall compliance and impaired ventricular relaxation restrict chamber filling and size in non-survivors, despite similar or potentially greater (see below) increases in ventricular filling pressures compared to survivors.

**Figure 3:**
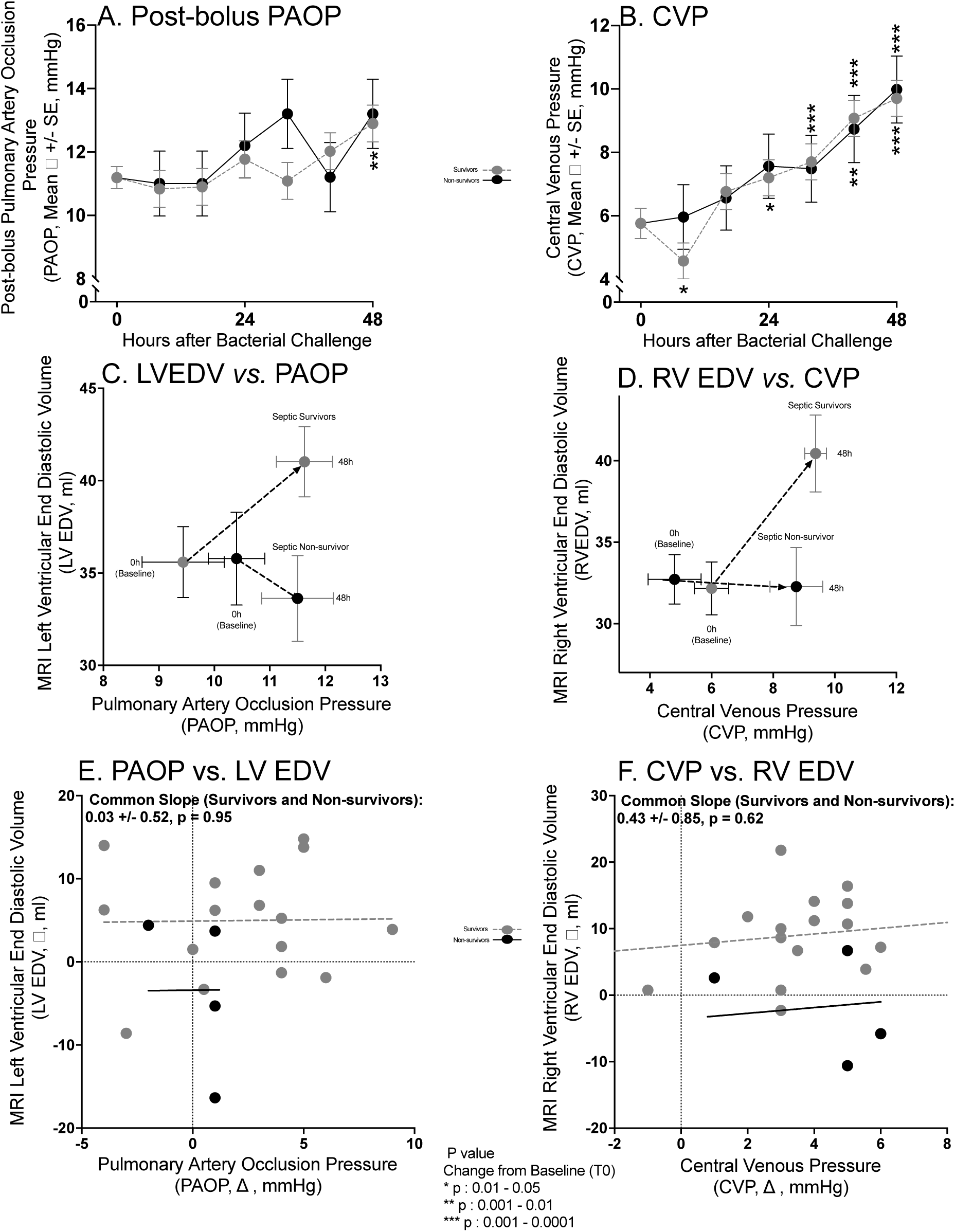
Loading conditions in septic survivors *vs*. non-survivors. The format is similar to Figure 2 except now serial mean post-fluid bolus PAOPs are shown in Panel A and serial mean CVP in Panel B. Ventricular Compliance plots of pressure *vs*. volume are shown for the left-sided circulation (Mean PAOP *vs*. LVEDV) in Panel C and the right-sided circulation (Mean CVP *vs*. Mean RVEDV) in Panel D. Correlations of changes in PAOP *vs*. LVEDV in Panel E and changes in CVP *vs*. RVEDV in Panel F.

Since all animals received similar maintenance fluids, the number of fluid boluses received was examined next (Figure 4, Panel A). Septic survivors compared to non-survivors received significantly more fluid boluses in the first 24h (Figure 4, Panel B). Accordingly, the PAOP fell below 10mmHg significantly more frequently in survivors than non-survivors, triggering the fluid bolus algorithm more and therefore increasing the number of boluses. From 26-42h, the number of fluid boluses received comparing septic survivors and non-survivors were not significantly different. More importantly, overall fluid balance did not differ between septic survivors and non-survivors from 0 to 48h (Figure 4, Panel A).

**Figure 4:**
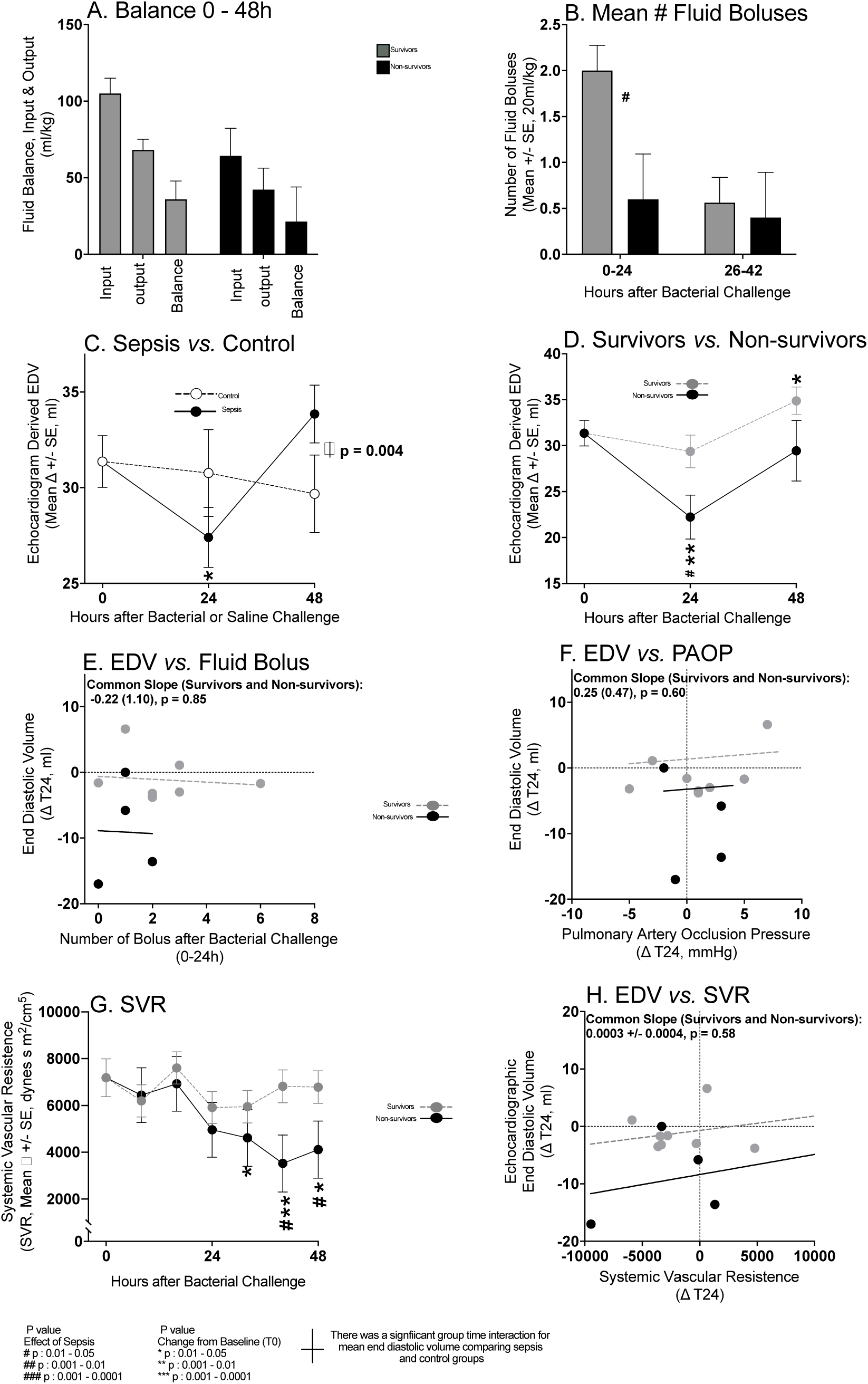
Loading conditions day one of sepsis. The format for Panel A, B, D to H is similar to Figure 2 and 3. The format for Panel C is similar to Figure 1. EDV values are ascertained by TTE.

We next determined whether fluid boluses in survivors compared to non-survivors in the first crucial 24h might explain observed changes LVEDV at 24h. As CMRs at 24h were not obtained, we instead evaluated TTEs changes. Serial TTE derived mean LVEDVs varied significantly more in septic animals from 0-24h and 24-48h compared to non-septic controls (Figure 4, Panel C, group/time interaction, p=0.004). In non-septic controls, there were no significant changes from baseline in mean LVEDV throughout (Panel C). Unexpectedly, in the crucial first 24h, the mean LVEDV in septic animals significantly decreased from baseline (Figure 4 Panel C) and this decrease was significantly greater in non-survivors (Figure 4 Panel D).

There were no hemodynamic changes in septic survivors and non-survivors to explain the 0-24h decrease in mean End Diastolic Volume (EDV) as follows: no significant mean decreases from baseline, or differences comparing groups for mean PAOP from 0-24h (Figure 3 Panel A and e-supplementary Figure 2, Panel A); no significant relationship in either group between the decreases in LVEDV and the number of fluids boluses received 0-24h (Figure 4, Panel E) or the changes in PAOP (Figure 4, Panel F); no significant changes in serial mean SVR 0-24h (Figure 4, Panel G) or marked increases in mean HR 0-24h (e-supplementary Figure 2, Panel D); no significant relationship between the decreases in LVEDV and the changes in SVR 0-24h (Figure 4, panel H) and HR 0-24h (e-supplementary, Figure 2, Panel E). From baseline to 24h, mean HRs in non-survivors were, on average, 110-120 beats per minute and even lower in survivors (e-supplementary Figure 2, Panel D). This range of HRs is unlikely to hinder cardiac filling and decrease EDV. Instead, it is likely a compensatory response with higher HRs in non-survivors to preserve cardiac output as the ventricle is unable to dilate and increase filling to maintain SV. The decreased chamber size in non-survivors *vs.* survivors was associated with a significantly greater decreased mean SV from 0-48h (P=0.05, e-supplementary Figure 2, Panel F). This loss of ventricular volume in the first 24h not attributable to decreases in filling pressures, or changes in afterload is most consistent with a sepsis-induced “restrictive-like” cardiac physiology.

Next, different aspects of LV wall structure were examined by CMR to better understand the physiologic underpinnings of cardiac dysfunction and ventricular volume changes during sepsis. As seen in e-supplementary Figure 3, Panel A (48h) and Figure 5, Panel A (96h), most of the changes in endocardial volume (i.e., LVEDV) *vs.* the epicardium volume at 48h and 96h fall below the identity line. This indicates that there are greater increases in the endocardial surface relative to the epicardial surface or that the LVEDV is changing more than the epicardial volume. Therefore, the distance between the two walls is getting smaller such that the LV wall is thinning. In controls, the volumes derived from tracing the epicardial and endocardial walls have values that appear closer to the identity line, indicating the LV wall is not thinning (e-supplementary Figure 3 Panel B). This wall thinning in septic animals was associated with significant decreases in wall mass from 0-96h (e-supplementary Figure 3 Panel C) which was likewise not seen in controls. The decreased wall mass comparing septic to control animals approached statistical significance (P=0.055).

**Figure 5:**
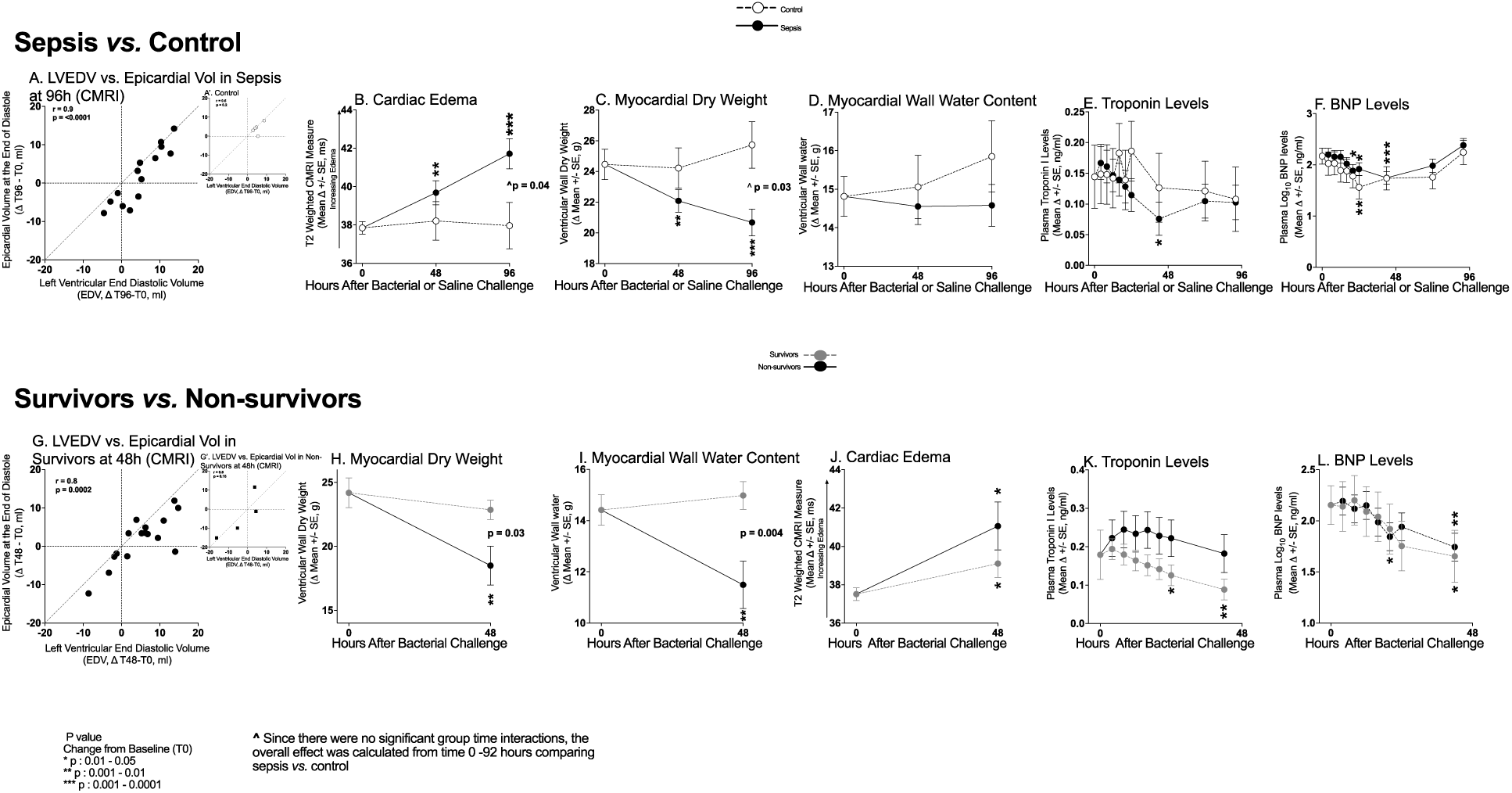
Serial Changes in Ventricular Walls during Sepsis. The format for Panel A – F is similar to Figure 1. The format for Panel G to L is similar to Figure 2 and 3.

Simultaneously, water per unit area of LV wall (i.e., edema) from 0-48h (∼2-3%) and 0-96h (∼4-6%) significantly increased in septic animals compared to controls (Figure 5, Panel B). Using these CMR measurements, LV dry (Figure 5, Panel C) and wet weight (Figure 5, Panel D) were each calculated (see e-supplementary Statistical Methods). In septic animals, from 0-96h, there was a significant loss of LV dry weight (p=0.03) but not water content. Pertinent blood work related to the myocardial walls showed that troponin I and BNP levels were not significantly elevated nor different from controls throughout (Figure 5, Panel E & F respectively). Thus, there was no evidence of fluid overload causing atrial stretch and LV dilatation in survivors nor was there evidence of ischemia causing myocyte loss during sepsis to explain the decrease in LV wall mass.

The changes in endocardial *vs.* the epicardial volumes at 48h in septic survivors also fell below the identity line, consistent with wall thinning (Figure 5, Panel G). Associated with this wall thinning, ventricular wall mass decreased significantly more in non-survivors compared to survivors at 48h (e-supplementary Figure 3 panel D). Non-survivors also lost more LV (dry mass) tissue (-5.69 +/- 1.52 g *vs.* -1.33 +/-0.76 g, p=0.03) (Figure 5, Panel H), more water content (-2.92 +/-0.92 g *vs.* + 0.56 +/- 0.54, g p=0.004) (Figure 5, Panel I), and had nominally greater increases in edema in the LV wall (Figure 6, Panel J). In non-survivors the mean troponin I (Figure 6 Panel K) and BNP (Panel L) levels were also not significantly elevated from baseline nor significantly different than survivors throughout.

**Figure 6:**
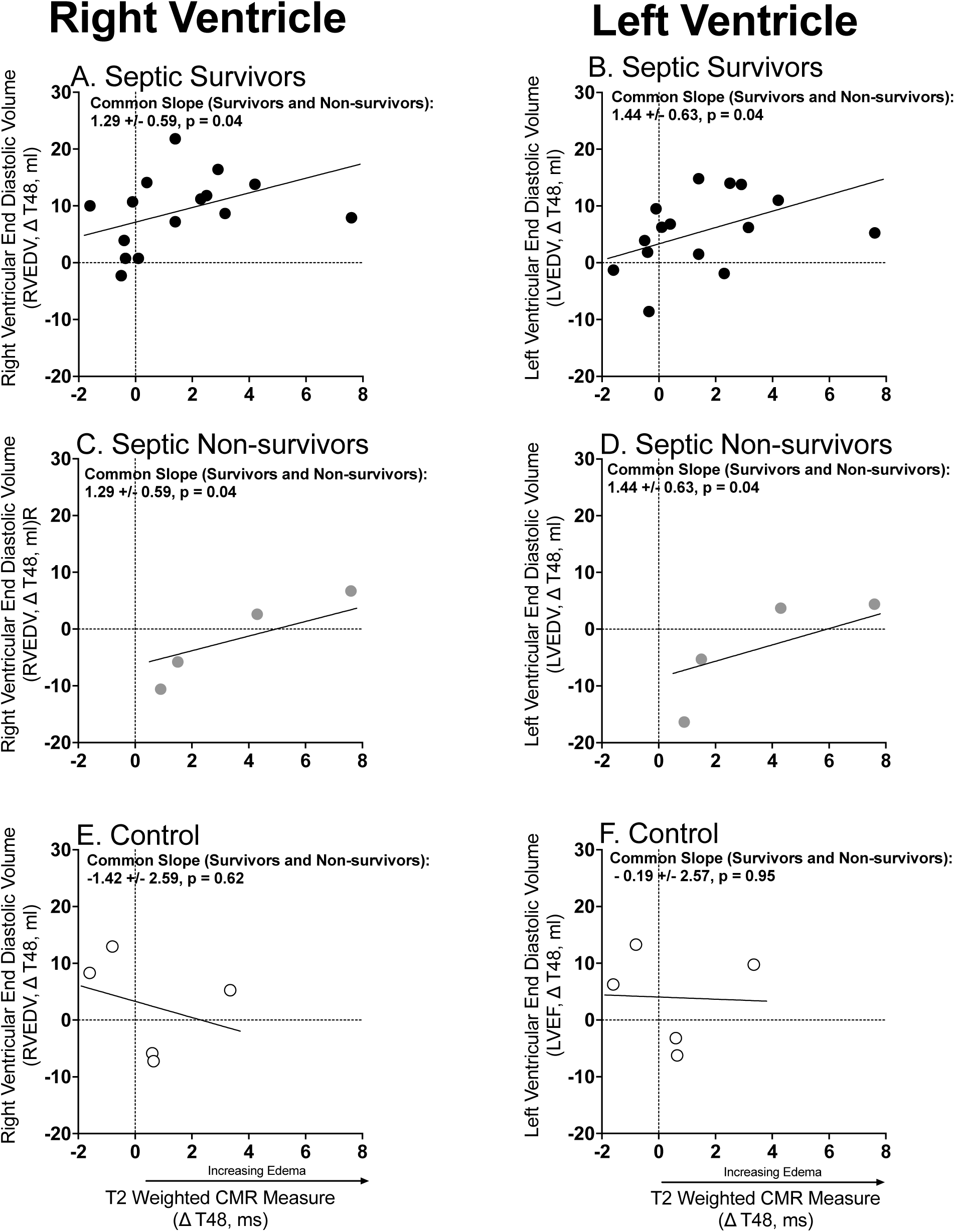
Association between MRI Derived Cardiac Edema and L and R End Diastolic Volumes. The format is like Figures 2 and 3.

Lastly, as ventricular wall water content increased in the septic survivors (Figure 6, Panel A and B) and non-survivors (Panel B and D), RVEDV and LVEDV also increased. No such relationship was seen among non-septic controls (Figure 6 Panels E&F). Thus, as the LV wall thins due predominantly to tissue loss rather than water, the LV chamber size, and overall total percent of water content increases. Sepsis is defined by initial ventricular volume decreases followed by chamber dilation, LV wall thinning with tissue loss and a relative increase in wall edema.

The morphological analysis of ventricular tissue collected at 62h from six septic animals confirmed previous cardiac histopathology and EM results.^13^ On histology, the LV had focal interstitial edema (Figure 7, Panel A) but no findings consistent with inflammation, microvascular occlusion, nor myocyte necrosis. EM of myocytes showed mild myofilament loss and intra-cellular edema (Figure 7, panel B) but no necrosis or other degeneration. The main EM findings were of non-occlusive capillary endothelial cell injury as evidenced by increased edema. Specifically, the endothelial cells making up the capillaries showed a mix of findings with mild to moderate edema and no changes in many endothelial cells. (Figure 7, Panel C). Although these results are qualitatively like prior histological and EM findings at 48h after bacterial challenge in a similar model;^13^ quantitatively, the endothelial and myocyte changes and edema appeared milder at this later time point (62h).

**Figure 7:**
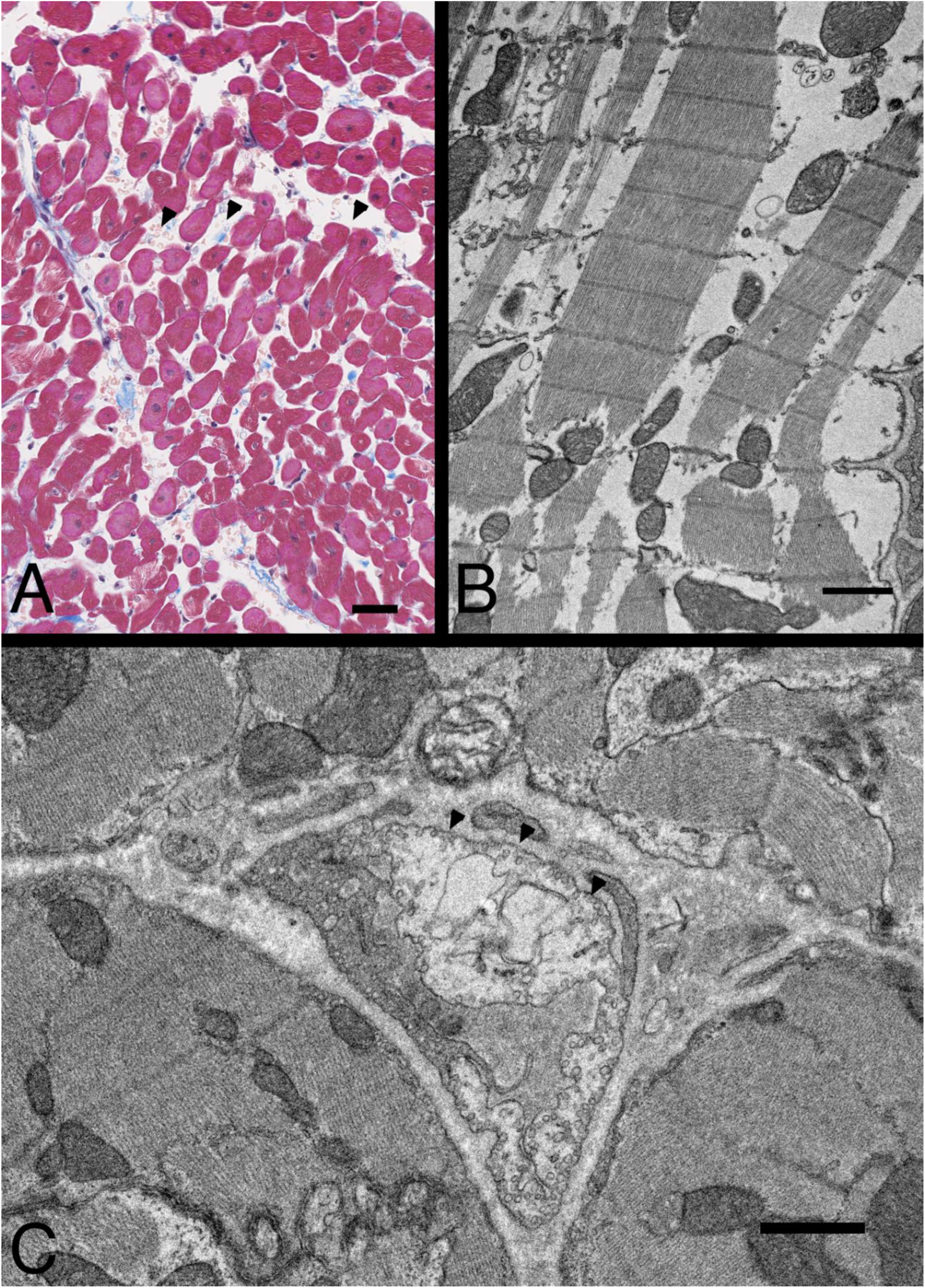
Representative images of Histology and electron microscopy. A. Micrograph of LV wall stained with Masson trichrome, shows myocardial interstitial edema (arrow heads). Bar, 25 μm. B. Myocyte with myofilament loss and intracellular edema. C. Partially damaged capillary endothelial cells. Note that the endothelial cell right side clearly shows edema (arrowheads), while the endothelial cell on the left side shows normal density. Bars, 1 μm.

There were isolated significant findings comparing mean serial values for septic animals *vs.* non septic controls and septic survivors *vs.* non-survivors for serum cytokines, chemistries, complete blood count, electrolytes, and arterial blood gas parameters that do not explain the observed cardiac findings or changes in EDV associated with outcome during sepsis. These results (e-supplementary Figures 1-13) and times of deaths of non-survivors (e-supplementary Table 1) are available in an e-supplementary Results.

## Discussion

During sepsis and into a relatively rapid recovery phase, there is an increase in edema and loss of dry tissue mass within the LV wall. During the first 24h of sepsis, there are decreases in ventricular volumes consistent with injury associated with the development of wall edema and diastolic dysfunction. Later, between 24 to 48h, ventricular volumes increase and dry mass decreases along with ventricular wall thinning. The extension and the acceleration of these changes in survivors out to 96h into the early recovery period suggests that they may represent the physiologic manifestations of sepsis resolution and cardiac repair.

These same ventricular volumetric chamber changes have been reported previously in this model.^6, 7^ However, this was thought to be the sequela of an inflammatory response that caused vasodilation and low filling pressures requiring fluid resuscitation and potent vasopressors that subsequently affected ventricular volumes. This study challenges those assumptions. Throughout, loading conditions were tightly controlled, exogenous catecholamines were not used, and cardiac filling pressures were optimized using invasive hemodynamic monitoring. Despite these measures, the same previously documented changes in ventricular chamber size and differences in survival were again seen. Here, preload, afterload or HR did not explain the evolution of ventricular volume changes that occur during septic shock. This indicates that septic shock must directly alter the structure and function of ventricular walls to produce these sepsis-induced changes in RV and LV chamber size.

The first cardiac abnormality observed after bacterial challenge was a deterioration in ventricular compliance where non-survivors developed a restrictive ventricular physiology. This “restrictive-like” cardiomyopathy was associated with decreasing chamber size without an associated decrease in cardiac filling pressures. Loading conditions alone were insufficient to explain this as all animals were fluid resuscitated to appropriate filling pressures. In fact, non-survivors required less fluid resuscitation during the first 24h after bacterial challenge. Non-survivor PAOPs fell below 10mmHg less frequently and required fewer fluid boluses. Therefore, in the first 24h, non-survivors have smaller ventricular chambers with fewer fluid boluses than survivors. As such, non-survivors compared to survivors had serially higher filling pressures and smaller chamber sizes, a pattern consistent with decreased ventricular compliance. Diastolic dysfunction with inadequate ventricular relaxation appeared to limit cardiac filling and impair forward flow particularly among non-survivors. Biventricular systolic and diastolic dysfunction seen during sepsis is a well-recognized entity in patients with heart failure and has been associated with a very poor prognosis.^17, 18^

After bacterial challenge, the major abnormality found by both histology and CMR was ventricular wall edema. We found histologic and biochemical evidence that myocytes were relatively preserved. EMs demonstrated only minimal focal myofilament loss in myocytes and no obvious myofilament remodeling or significant structural derangements with mostly mild edema and no inflammatory cell infiltrates. Troponin I levels were not elevated throughout the study, arguing against cell death or tissue integrity loss due to ischemia. Plasma BNP levels were also not elevated, suggesting fluid overload did not induce myocyte stretch with resulting increases in biventricular EDV and ESV. We found no other obvious histopathological cardiac abnormalities that explained the observed increases in RVEDV and LVEDV or the reversible profound falls in LVEF after bacterial challenge.

Myocardial edema has been strongly associated with cardiac dysfunction in animal models of acute cardiac disease.^19^ Small increases in myocardial edema was found to compromise cardiac function in both acute and chronic models of hypertension in canines,^20^ with other models reporting a reduction of cardiac function by 40%.^19, 21^ In a canine coronary venous hypertension model, as myocardial edema increased, there were elevations in LV chamber stiffness and reductions in ED*Vs.*^22^ In septic mice, decreases in LVEF were associated with myocardial vascular injury and leakage,^23^ resulting in increased edema and cardiac dysfunction.

Cardiac edema has been reported in cardiac human sepsis studies employing cardiac EM and CMR. This agrees with our cardiac EM and CMR data and previously reported preclinical cardiac EM data.^13^ An observational study of 15 septic patients showed increased myocardial T2-times and decreases in LVEF within 48h after ICU admission.^24^ Three septic patients in another study showed increased T2-signal and histological evidence of “striking” interstitial edema.^25^ An autopsy report described two septic subjects with cardiac dysfunction and myocardial edema.^26^ Therefore, edema is a plausible antecedent mechanism of cardiac dysfunction during sepsis given 1) the rapidity of cardiac injury [chamber size changes and drop in LVEF and RVEF] and speedy reversal over days; 2) the findings of edema on CMR and histology in septic humans and animals; 3) the degree of edema found and the consistency across the literature of its association with reversible cardiac dysfunction; and, lastly, 4) no other obvious major histopathological injury was found to explain the EDV and LVEF findings. As this controlled study shows a crucial link between cardiac edema and dysfunction during sepsis, a well-documented feature in the literature by CMR and EM, we suggest the following acronym: <u>S</u>eptic <u>C</u>ardiac <u>E</u>dema <u>R</u>elated <u>R</u>eversible Injury (SCERRI).

Initially, decreased ventricular chamber size at 24h was followed by a marked increase from 24-48h in all septic animals. We also found a significant strong positive correlation during this time between increases in EDV and LV wall edema among septic animals. This correlation reinforces the concept that EDV increases are closely related to the loss of ventricular dry mass. LV wall edema increased because LV wall dry weight decreased without a loss of water, which therefore led to a relative increase in the water content of the LV wall. Simultaneously, as mass loss thinned the LV wall, a corresponding increase was observed in EDV. Collectively, these CMR-measured effects were reflected in the positive correlation between EDV and edema in both survivors and non-survivors. The loss of mass continued from the onset of infection throughout recovery of cardiac function, with an approximate 15% decrease from baseline in LV wall dry weight by 96h. Because mass loss continued during LVEF recovery, we speculate that this is part of a reparative process in survivors. Non-survivors presumably have greater, non-recoverable cardiac injury and were found to have significantly greater LV dry mass loss at an early timepoint (48h), just prior to death. If dry mass loss is a manifestation of a repair process, it started early and was stronger in sicker animals, but was not able to overcome an otherwise unrevivable episode of septic shock.

Surprisingly, we found that cardiac volumetric changes (increased LVEDV and RVEDV) rather than functional ones (i.e., LVEF, strain, ventricular-arterial coupling) correlated with outcome. Since greater LV chamber dilation and larger LVEF decreases have been associated with better outcomes in humans sepsis,^27^ we evaluated the human cardiac sepsis literature to assess its compatibility with our results [e-supplementary Meta analysis (Methods, e-supplementary Figure 14 and Results e-supplementary Tables Two - Four)]. Overall, our animal data reported here, and the human literature agree: LV chamber dilation is associated with sepsis survival, but not changes in LVEF.

Evidence for myocyte dropout, necrosis, or stretch was not found biochemically or histopathologically in our canine model of septic shock. Therefore, our finding of ventricular dry mass loss must have occurred due to loss of intracellular or extracellular constituents or destruction of some other abundant, non-myocyte cardiac cell type. Cardiac endothelial cells constitute ∼65% of non-myocyte heart cells and have an integral role in cardiac remodeling and regeneration, and have been implicated in the modulation of myocytes contractility.^28,29^ Sepsis-induced dysfunction of this dynamic barrier through inflammatory injury with disruption of crucial intracellular signaling and cell-cell crosstalk can lead to increased vascular permeability and precipitate significant cardiac dysfunction.^30^ Endothelial disruption during sepsis has been well characterized in other organs. Liu *et al.* showed in murine lungs that inflammatory injury by LPS was associated with a loss of pulmonary endothelial cells which progressively recovered over the 7 days.^31^ This is consistent with the time frame of dry weight loss and recovery of cardiac function that we observed in survivors. This disrupted vascular endothelium and potential loss of endothelial cells that improves over 5-7 days provides a potential mechanism for cardiac edema and loss of ventricular mass in sepsis observed in our study. During sepsis, we observed some focal myofilament degradation on EM. Autophagy is a normal quality control process where eukaryotic cells form autolysosomes to degrade and remove damaged molecules and organelles. In murine endotoxin models, the activation of autophagy has been shown to reduced cardiac dysfunction.^32, 33^ Notably, autophagy with removal and replacement of injured or dysfunctional cell components has been found to be protective in several models of acute cardiac injury.^34–36^ The loss of extracellular matrix and matricellular proteins involved in injury and repair could also contribute to this tissue loss.^37^

There are limitations to this study. CMR scans were only obtained at baseline, 48h and 96h, leaving notable gaps in the timeline of sepsis-induced cardiac injury and dysfunction. However, TTEs were obtained daily which bridged these gaps and allowed us to follow some aspects of cardiac function. Secondly, different types, doses, and sites of bacteria could potentially result in different findings; however, our well-established animal model has demonstrated that the pattern of changes in LVEDV and EF is independent of these factors. Third, while echocardiographic diastology measures were collected there are no well-validated guidelines in canines, and the values obtained differ substantially from the range of human echocardiographic reference standards.

## Conclusion

This study provides evidence that changes in preload, afterload, or heart rate cannot explain changes in biventricular chamber size over time in septic survivors compared to non-survivors. In the absence of catecholamine administration and despite optimizing preload, non-survivors developed from 0-24h, a worsening restrictive physiology with less dilation compared to survivors despite similar Ejection Fractions and higher PAOPs. An increase in cardiac edema seen by CMR and histology was associated with cardiac injury and dysfunction. Furthermore, sepsis-induced loss of ventricular wall dry mass over 96h extended into the recovery of biventricular ejection fractions, indicating that it may be a reparative process rather than ongoing injury. This loss of mass in part can explain sepsis-induced increases in EDV from 24 to 48h. The loss of endothelial cells, extracellular matrix, and/or the autophagy and replacement of damaged endothelial and myocyte molecules and organelles may explain this loss of ventricular mass and thereby account for wall thinning during septic shock. The findings are important in that such changes are associated with outcome and therefore warrant further investigation. Finally, this is the first controlled CMR sepsis study to demonstrate that ventricular wall edema is a critical element of sepsis pathophysiology and dry mass loss may be part of the reparative process.

## Supporting information

Supplemental methods, results, figures, tables

## Non-standard Abbreviations and Acronyms

ABGs: Arterial Blood Gas
BNP: Brain Naturietic Peptide
CBCs: Complete Blood Count
CMR: Cardiac Magnetic Resonance
CVP: Central Venous Pressure
EDV: End Diastolic Volume
EM: Electronic Microscopy
HR: Heart Rate
LV: Left Ventricular
LVEDV: Left Ventricular End Diastolic Volume
LVEF: Left Ventricular Ejection Fraction
MAP: Mean Arterial Pressure
PBS: Phosphate Buffer Solution
PACs: Pulmonary Artery catheters
PAOP: Pulmonary Artery Occlusion Pressure
PAP: Pulmonary Artery Pressure
PVR: Pulmonary Vascular Resistence
RVEDV: Right Ventricular End Diastolic Volume
RNCA: RadioNucleotide Cine Angiocardiogram
RVEF: Right Ventricular Ejection Fraction
RV-PA: Right Ventricular – Pulmonary Artery
RVESV: Right Ventricular End Systolic Volume
RVSV: Right Ventricular Stroke Volume
SCERRI: Septic Cardiac Edema Related Reversible Injury
SV: Stroke Volume
SVR: Systemic Vascular Resistance
TTE: Transthoracic Echocardiogram

## SUPPLEMENTAL MATERIAL

Supplemental Methods

Supplemental Results

Supplemental Meta-analysis

Figure S1-S14

Tables S1–S4

References 38–57

