## Supplemental methods, results, figures, tables for "Cardiac Magnetic Resonance Studies in a Large Animal Model that Simulates the Cardiac Abnormalities of Human Septic Shock"

### eSupplement

#### e-supplementary Methods

##### Transthoracic Echocardiography

Daily serial transthoracic echocardiograms were performed using standard techniques for all animals from baseline until 96 h after bacterial challenge. Echocardiographic examination was performed using **a** 2.5-7.0 MHz transducer and simultaneous ECG recording (VIVID-IQ, General Electric Medical System, Waukesha, Wisconsin).

Animals received their echocardiogram in the right lateral and supine recumbency position. All images and measurements were collected from standard views and digitally stored for offline analysis. LVEF was calculated using Simpson's method using readings of the apical 4 chamber and 2 chamber bi-plan views or visual eyeball LVEF estimation.

LV diastolic function was assessed by trans mitral flow velocity (E and A) and E/A was calculated. Lateral and septal mitral annular flow velocities (e') were measured and E/e' lateral and septal were calculated. All measurements were calculated by an experienced technician and reviewed by a senior echocardiologist to avoid inter-observer bias.

##### Cardiac MR

Each animal was transported to the scanner sedated, mechanically ventilated, and continuously monitored by a technician or clinician. A 3 Tesla MRI scanner (Philips Healthcare) acquired cardiac MRIs for animals at baseline (T0), 42 hours after challenge, and at the end of the study (96h). Electrocardiogram-gated steady state free-precision cine and T2 images were acquired in mid-ventricular short axis and assessed for average plane values. Epicardial and endocardial contours were drawn on the short-axis slices at end-diastole and end-systole. For contrast enhancement, animals were given gadobutrol 0.1 mmol/kg (Bayer Healthcare). Throughout the scan, animals were sedated, ventilated, and continuously monitored by technicians and or clinicians.

All measurements for the CMRs were conducted using dedicated analysis software (NEOSOFT suiteHEART) by appropriately trained clinicians and further confirmed by an experienced independent clinician. Papillary muscles were included in the volumetric quantification of the left ventricle.

**Electron Microscopy and Histology**

Histology and electron microscopic analysis were performed as previously described.[13] Briefly, the left ventricle anterior wall tissues were fixed and embedded in paraffin. Sections were then stained with HE and Masson trichrome. The slides were scanned with NDP digital slide scanner (Hamamatsu) for histology evaluations. For Electron Microscopy (EM), the cardiac tissues were dissected from the left ventricle anterior portion and brief fixed with 4% formaldehyde and 1% glutaraldehyde then sliced to 200 m slices with vibratome, further fixed overnight, followed by post fixation with 1% $OsO_4$ and embedded in Epoxy. Ultrathin sections were stained with lead and uranyl and images were taken from JEOL-1400 electron microscope.

**Animal Inclusion Criteria**

Animals were pooled from two original experiments, the first assessed the effects of catecholamines on myocardial function during sepsis (n = 14) and non septic controls (n = 12), and the second assessed a bacterial dose response on the myocardial function in sepsis (n = 14). Of these total 40 animals, 27 animals did not receive catecholamines and were included in a secondary analysis in this manuscript. Six additional septic animals were employed for cardiac histology and EM.

**Statistical Methods**

Using CMR measurements, we calculated approximate “water content” as mass*edema (in %), and “dry weight” as mass – water content. Unless noted otherwise, data was analyzed using linear mixed models to account for repeated measures and plotted as model estimate (+/- SE). We first tested the group-time interaction. If the interaction term is significant, groups are compared at each time point; otherwise, group comparisons are based on the main effects. Standard residual diagnostics were used to check model assumptions. All *p*-values are two-sided and considered significant if $p \leq 0.05$. For some variables, logarithm transformation was used when necessary. Random-effect models were used to estimate the overall mean difference (MD) or standardized mean difference (SMD) between groups. Heterogeneity among studies was assessed using the Q statistic and $I^2$ value. Meta-analyses were conducted using *R* (version 4.2.2) and packages *meta* (version 6.2-0)[38] and *metaphor* (version 3.8-1)[39]. All other statistical analysis was performed using SAS version 9.4 (Cary, NC) and figure creation using GraphPad Prism 9.

**Supplementary Laboratory Results**

**Cytokines (IL-6, -8 -12 and -10, interferon- γ, von Willebrand factor, Tumor necrosis factor- α, p-selectin, and Monocyte Chemoattractant Protein)**

*Sepsis vs. controls*

In septic animals at 4, 6, 8, 12, 16, 20 and 24h after bacterial challenge, compared to baseline, there is a significant increase in mean IL-6 plasma levels (e-supplementary Figure 4, Panel A). Non-septic controls during this time had no significant changes from baseline in mean IL-6 plasma levels. Compared to controls, the mean IL-6 levels in septic animals were significantly elevated at 8, 12, 16, 20 and 24h after bacterial challenge (Panel A). The mean IL-8 levels at 6 and 8h after bacterial challenge (Panel B) and the mean IL-12 levels at 72 and 96h after bacterial challenge (Panel C) were significantly elevated compared to baseline. During this time, there were some significant decreases in mean IL-8 at later time points and no significant changes in mean IL-12 in controls throughout compared to baseline. Furthermore, there were no significant differences in the mean values of these two cytokines in controls compared to septic animals. In septic animal, mean IL-10 levels at 16 and 20 h (Panel D) were elevated and mean Interferon-γ levels (Panel E) at 6, 8, 12, 20 and 48h were depressed compared to non-septic controls. Additionally, septic animals did have significant depressions compared to baseline in the mean levels of Von Willebrand factor (VWF) (8 and 96h, Panel F), TNF- α (8,12,16,20, 48,72 and 96h, Panel G), and there were not significant differences during these times in mean values of the following cytokines (VWF, TNF- α and p-selectin) compared to controls. In septic animals at 4,6, and 8h after bacterial challenge, there were significant elevations of mean levels of Monocyte Chemoattractant Protein-1 compared to baseline (Panel I). In controls, there was no significant change in the mean value of this cytokine during this time compared to baseline.

Thus, the major cytokine elevated most profoundly in this sepsis model was IL-6; however, multiple other cytokines showed changes from baseline with either elevations or depressions that was not observed in controls.

*Septic Survivors vs non-Survivors*

There were no significant differences in the mean changes from baseline at 48h for any of the 9 cytokines measured (e-supplementary Figure 5 Panels A-I) between septic survivors and non-survivors. Of note, during this time 5 cytokines were nominally higher in survivors (IL-6, -10, Interferon- γ, von Willebrand factor, TNF- α). Two cytokines (IL- and -12) were nominally only minimally elevated over time in survivors and two of the nine Cytokines were remarkably similar over this time (p-selectin and Monocyte Chemoattractant Protein-1) in survivors and non-survivors. Overall, there were no remarkable increases in cytokine levels in non-survivors *vs.* survivors in this experimental sepsis model and if anything, survivors overall had higher cytokine levels not lower.

**Serum Chemistries, Complete Blood Count, Arterial Blood Gases, and Electrolytes**

*Sepsis vs. controls*

In septic animals, from 4 to 96 h after bacterial challenge, mean pH was significantly decreased from baseline at all time points measured. In non-septic controls, mean pH is only significantly decreased from baseline at 24 to 72h after bacterial challenge. However, there is overall from baseline to 96h a significant decrease in mean pH comparing septic and control animals (e-supplementary Figure 6**,** panel A all, p=ns). There were no significant changes from baseline in mean PCO2 in both septic and control animals throughout the study, as this parameter was maintained within a set range by protocolized mechanical ventilation (Panel B). Arterial mean PaO2 in septic animals was significantly lower *vs.* controls at 8 and 12h after bacterial challenge and not significantly different from controls at all other time points measured. This parameter was also artificially maintained by protocolized titration of inspired FiO2 on the mechanical ventilator (Panel C). Mean lactate levels were not significantly different in septic animals compared to controls throughout the duration of the study (Panel D).

There were no significant differences in change from baseline for mean Creatinine, BUN, Total Protein, Alanine Transaminase, and Albumin levels in septic *vs.* control animals throughout the study (e-supplementary Figure 7). In septic animas there were, at random time points, isolated significant differences compared to baseline (at levels of no clinical significance) in mean BUN (Panel B), BUN/Cr ratio (Panel C), Total protein (Panel D) and Albumin (Panel F). Septic animals had significant increases in mean glucose levels at 12, 16, 20, 24, 48, 72, and 96h compared to baseline. Mean glucose levels were normal in controls were significantly higher in septic animals throughout the study (e-supplementary Figure 8, Panel A). There were significant increases compared to baseline in mean potassium levels at 24, 48, 72, and 96h only but no significant difference from controls throughout. There were no other significant differences between septic and control animals in mean sodium (Panel C), and Chloride (Panel D) levels.

Septic animals had significantly lower mean WBC counts at 8, 12, 16, and 20h compared to controls (e-supplementary Figure 9**,** Panel A). There were no significant differences in mean Hemoglobin, Hematocrit, Lymphocyte, Eosinophil, and Platelet levels between septic animals and non-septic controls throughout the study (Panel B-F).

*Survivors vs. Non-survivors*

In survivors *vs.* non-survivors, there were no significant differences in mean pH (Panel A), PCO2 (Panel B), PaO2 (Panel C) and Lactate (Panel D) throughout the study (e-supplementary Figure 10). Non-survivors had significant increases compared to baseline in mean creatinine levels at 16, 20, and 24h. Mean creatinine levels were normal in survivors throughout the study but significantly lower compared to septic non-survivors (e-supplementary Figure 11, Panel A) at 16, 20, and 24h after bacterial challenge. In non-survivors, BUN levels were significantly elevated both compared to survivors (Panel B) and from baseline at 16, 20, 24, and 48h. There were no significant differences in mean BUN/Cr ratio, Total Protein, Alanine Transferase, and Albumin between survivors and non-survivors throughout the study (Panel C-F).

Mean glucose levels were not significantly different in survivors *vs.* non-survivors throughout the study (e-supplementary Figure 12, Panel A). Potassium levels were significantly increased from baseline in non-survivors at 24 and 48h (Panel B). Sodium (0 to 48h) and Potassium levels (0 to 48h) were significantly higher in non-survivors compared to survivors (Panel B and C, $p = 0.03$ and 0.02 respectively). There were no significant differences in chloride levels between survivors or non-survivors and no significant changes from baseline throughout the study (Panel D).

There were no significant differences in WBC, Hemoglobin, Hematocrit, Lymphocyte, Eosinophil, and Platelet levels between survivors and non-survivors of sepsis throughout this study (e-supplementary Figure 13, Panel A-F).

**e supplementary Literature Search for Meta-analysis**

A search of electronic databases PUBMED, EMBASE, and BIOSIS was performed to identify studies from 1980 until 02/22/2023. Any study investigating LVEF and EDV in septic survivors and non-survivors through ECHO, MUGA or CMR was included. Relevant titles were also identified by hand searching previous reviews on the topic. Duplicates were filtered through EndNote once the databases were combined. Studies that were excluded were pediatric populations, conference abstracts, reviews or editorials, non-sepsis related cardiomyopathies, or did not use the imaging modalities listed above (e-supplementary figure 14). Search terms that were used included cardiomyopathy OR "contractile dysfunction" OR "systolic failure" OR "diastolic failure" OR "impaired cardiac relaxation. These terms were combined with “ejection fraction" OR "diastolic volume" OR "diastolic volume index" OR ‘echo’ OR ‘echocardiography’, as well as the results from the combined set, 'sepsis' OR 'septic shock'. Results were screen further for imaging using, ‘mri’ OR ‘magnetic resonance imaging’; ‘rnca’ OR ‘radionuclide cineangiogra’’; ‘t2 weight’ OR ‘t2wi’ AND edema. /exp OR 'septic shock'.

**e-supplementary Results of Meta-analysis**

We found 20 septic studies since 1980 comparing LVEF in survivors and non-survivors[4, 16, 27, 40-56] (e-supplementary Figure 14) of which 12 reported LVEDV by outcomes.[4, 16, 46-51, 53, 54, 57] Across studies, at day one of sepsis, LVEFs varied widely (30.9 - 68.8%) (e-supplementary Table 2);

however, the mean difference in LVEF depression between survivors and non-survivors remained minimal across studies with overall mean difference of 1.0% (95% CI of -0.4 to 2.4%, p=0.15) with a low $I^2 = 5\%$. The LVEDV was reported multiple ways and, therefore, the standard mean difference (SMD) was calculated. The $I^2$ was 87% across the 12 studies, too high to justify combining data. One study caused the very high $I^2$; its exclusion profoundly reduced the $I^2$ and results in a highly significant summary finding (e-supplementary Table 3). In the 11 remaining studies, survivors had a larger LVEDV (SMD 0.36, $P \leq 0.0001$, $I^2 = 57\%$) (e-supplementary Table 4).

#### e-supplementary Figure 1: Loading Conditions and Afterload During Experimental Septic Shock

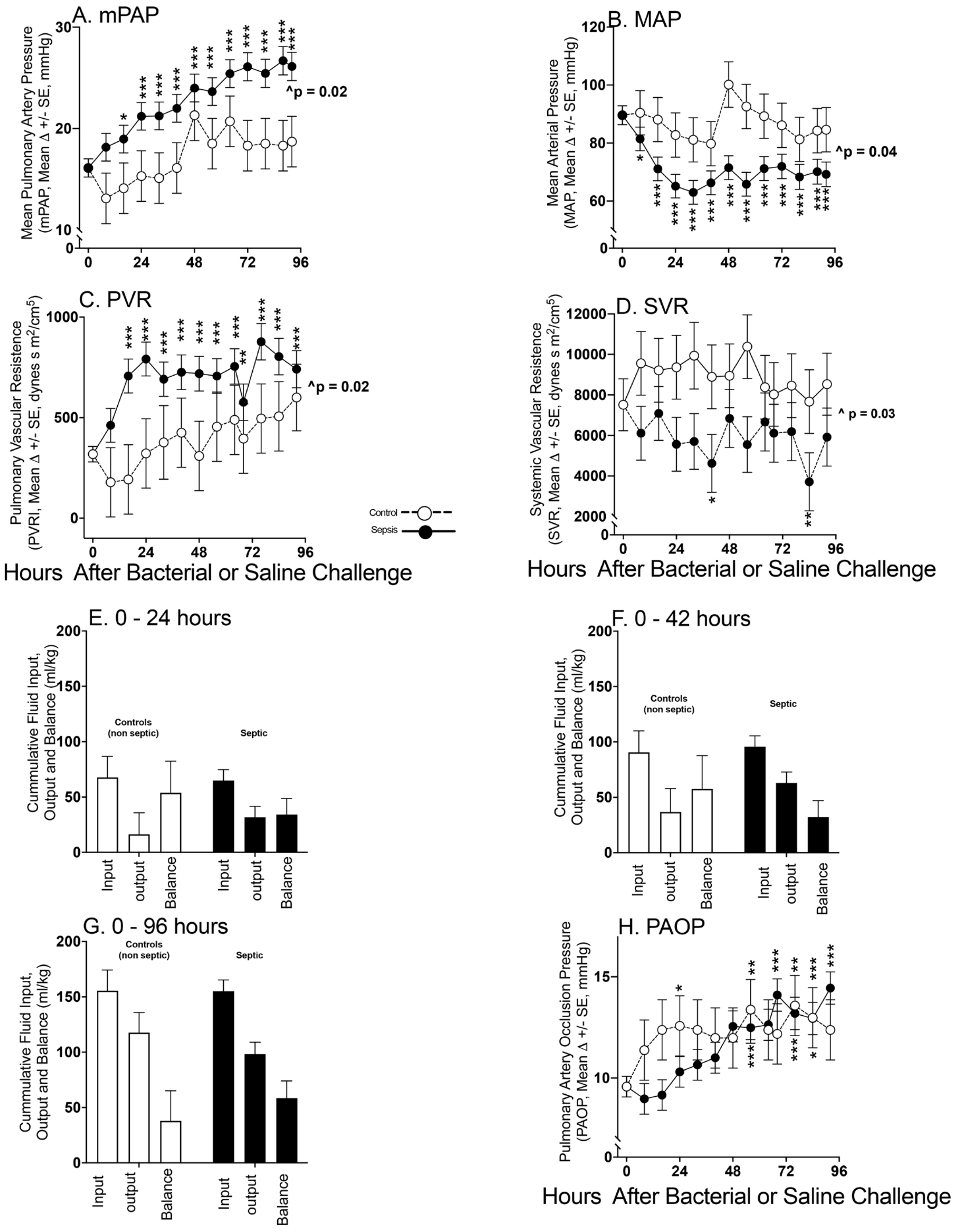


P value
Change from Baseline (T0)
* p : 0.01 - 0.05
** p : 0.001 - 0.01
*** p : 0.001 - 0.0001

^ Since there were no significant group time interactions, the overall effect was calculated from time 0 -92 hours comparing sepsis *vs.* control

**e-supplementary figure 1: The format is similar to Figure 1.**

**e-supplementary Figure 2:**

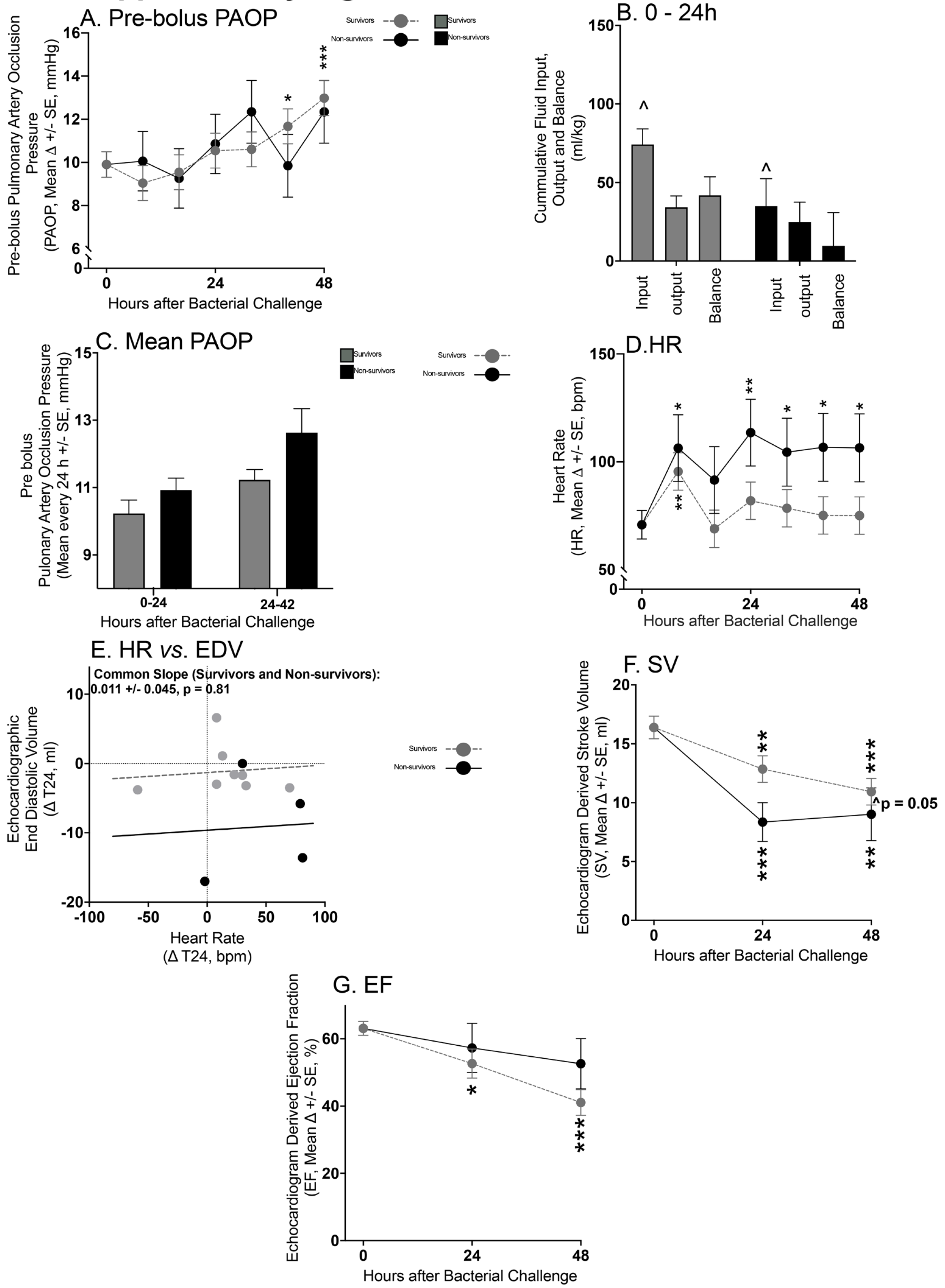


P value
Change from Baseline (T0)
* p : 0.01 - 0.05
** p : 0.001 - 0.01
*** p : 0.001 - 0.0001

^ p = 0.07 since there were no significant group time interactions, the overall effect was calculated from 0 - 48h comparing survivors *vs.* non -survivors. This was reported as it is borderline signifcant

**e-supplementary figure 2: The format is similar to Figures 1 and 2.**

#### e- supplementary Figure 3

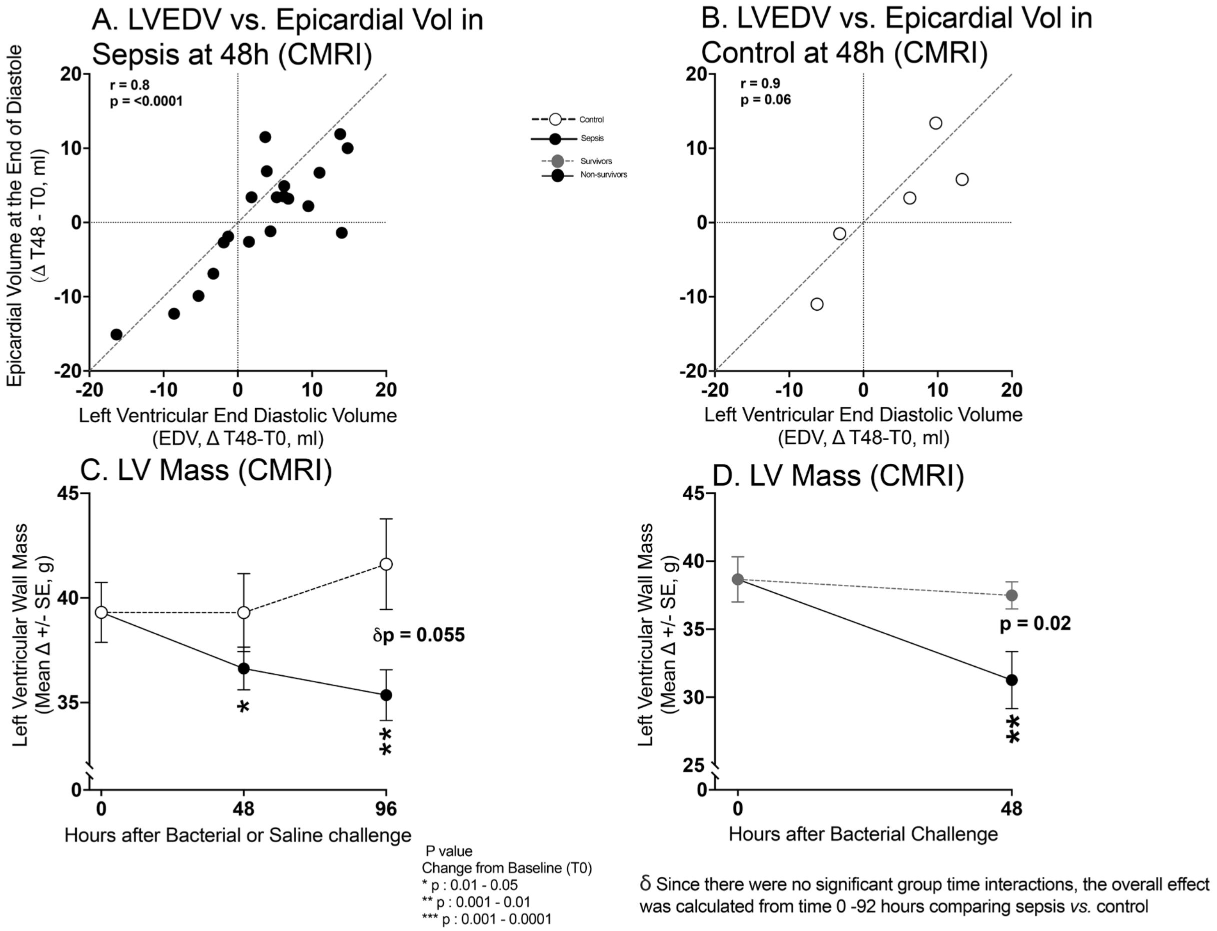


**e-supplementary Figure 3: For Panel A and B, the format is similar to Figures 5 Panel A. For Panel C, the format is similar to Figure 1. For Panel D, the format is similar to Figure 2.**

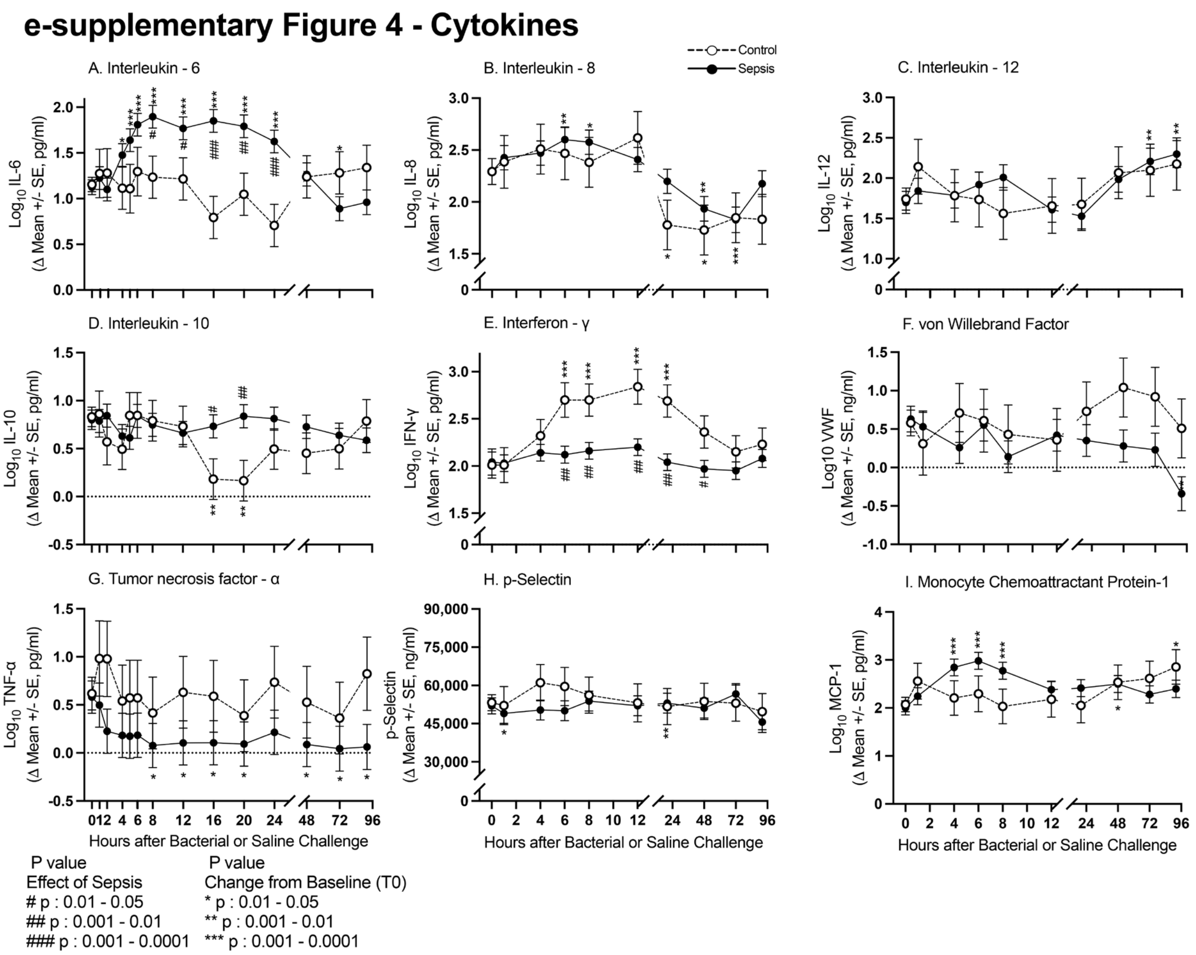


**e-supplementary figure 4: The format is similar to Figure 1.**

#### e-supplementary Figure 5 - Cytokines

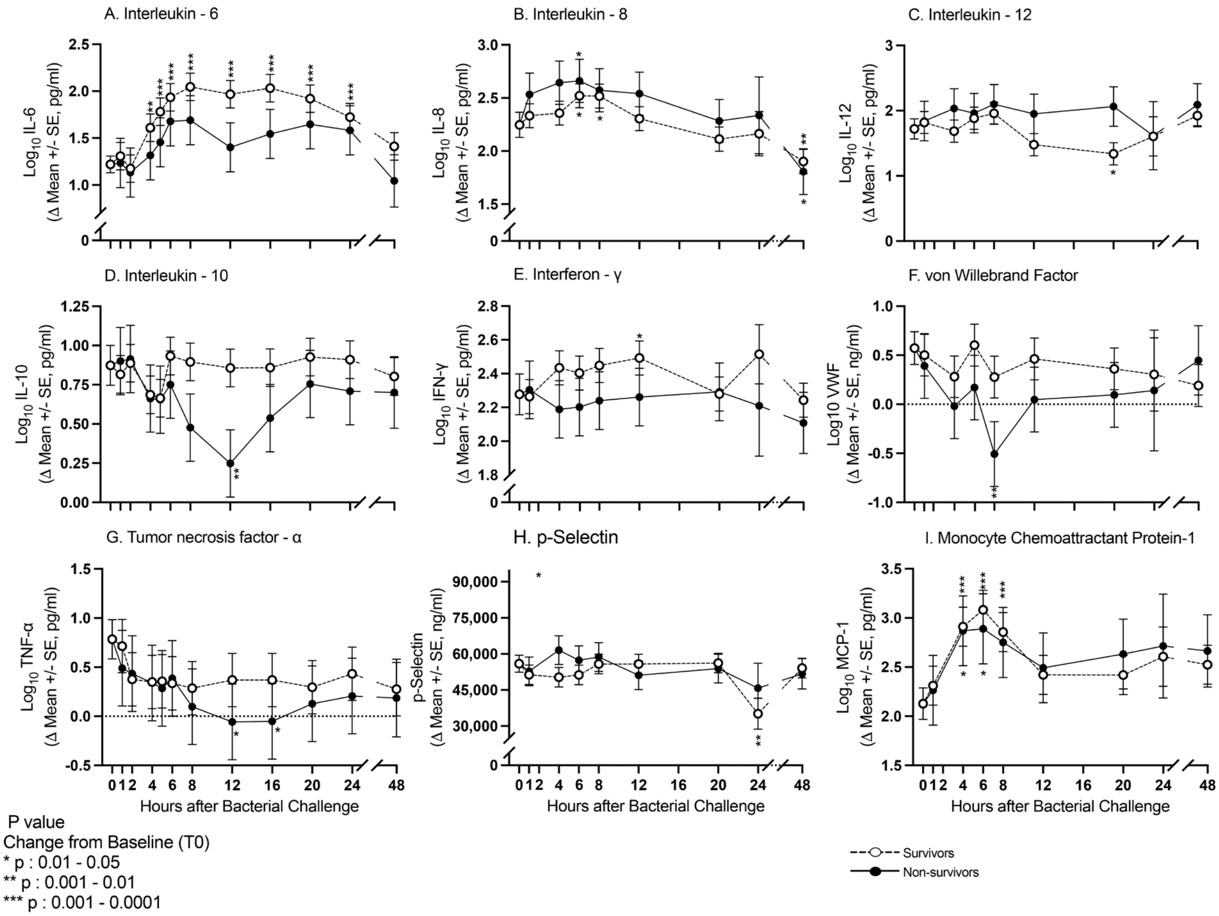


**e-supplementary figure 5: The format is similar to Figure 2**

#### e-supplementary Figure 6 - Arterial Blood Gass Measures

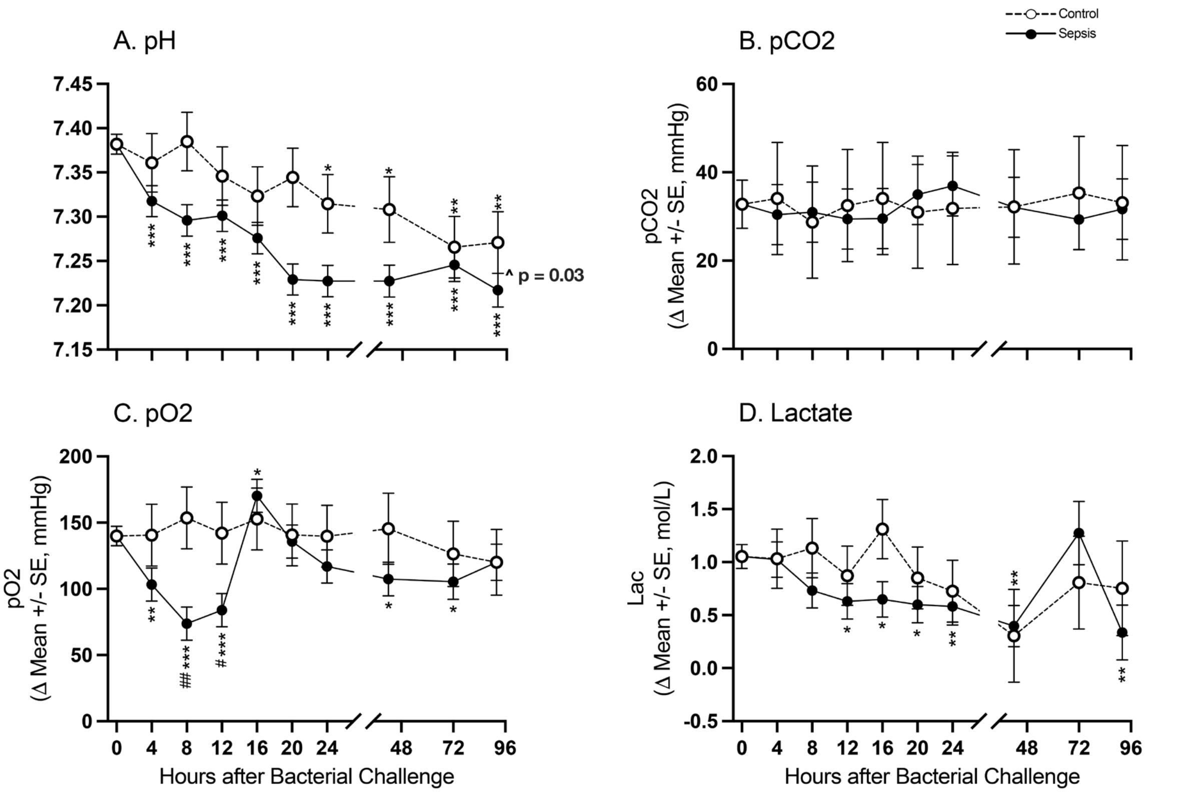


P value
Effect of Sepsis
# p : 0.01 - 0.05
## p : 0.001 - 0.01
### p : 0.001 - 0.0001

P value
Change from Baseline (T0)
* p : 0.01 - 0.05
** p : 0.001 - 0.01
*** p : 0.001 - 0.0001

^ Since there were no significant group time interactions, the overall effect was calculated from time 0 -92 hours comparing sepsis *vs.* control

**e-supplementary figure 6: The format is similar to Figure 1**

### e-supplementary Figure 7 - Serum Chemistries

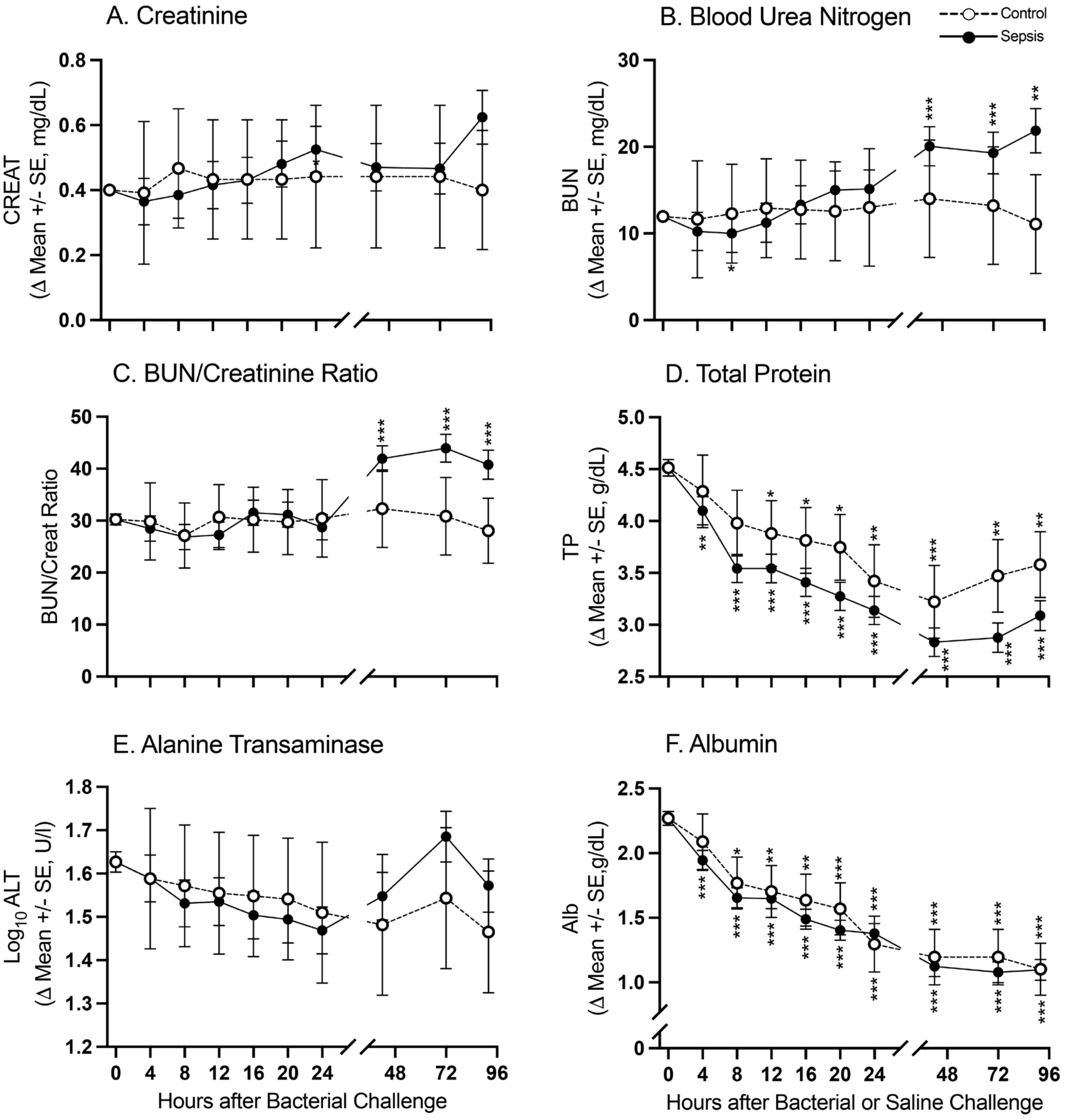


| P value Effect of Sepsis | P value Change from Baseline (T0) |
|---|---|
| # p : 0.01 - 0.05 | * p : 0.01 - 0.05 |
| ## p : 0.001 - 0.01 | ** p : 0.001 - 0.01 |
| ### p : 0.001 - 0.0001 | *** p : 0.001 - 0.0001 |

**e-supplementary figure 7: The format is similar to Figure 1**

#### e-supplementary Figure 8 - Serum Chemistries

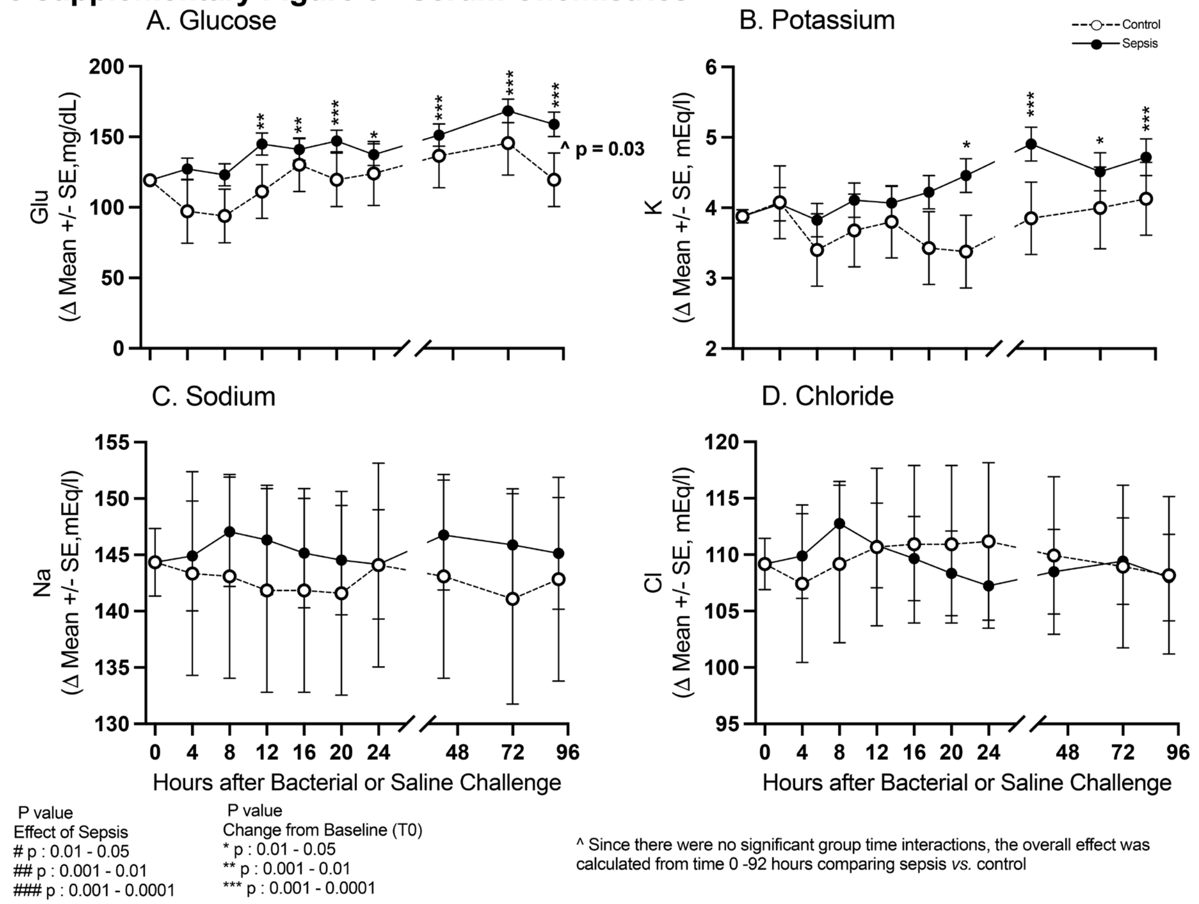


**e-supplementary figure 8: The format is similar to Figure 1**

### e-supplementary Figure 9 - Complete Blood Count

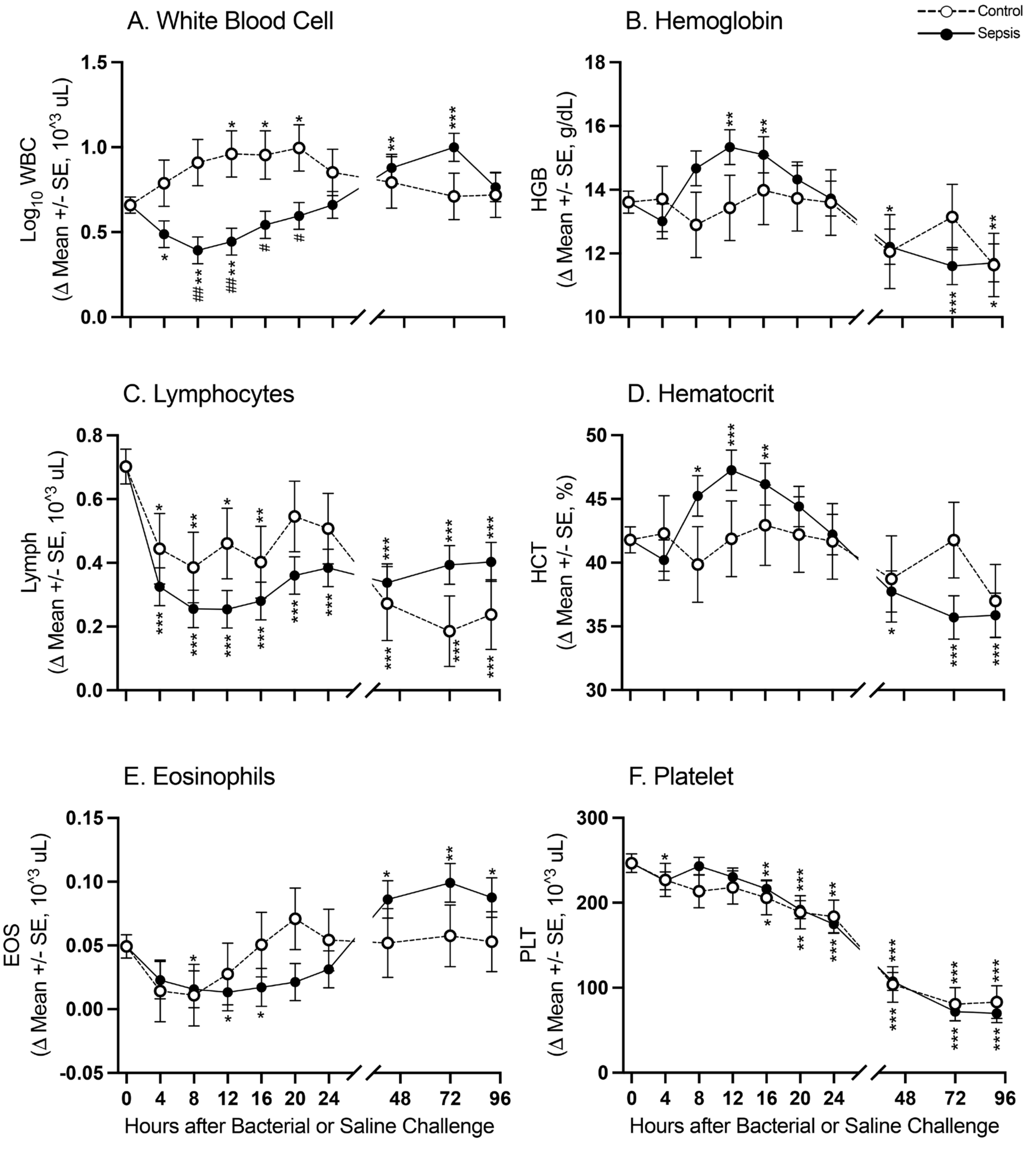


| P value | P value |
|---|---|
| Effect of Sepsis | Change from Baseline (T0) |
| # p : 0.01 - 0.05 | * p : 0.01 - 0.05 |
| ## p : 0.001 - 0.01 | ** p : 0.001 - 0.01 |
| ### p : 0.001 - 0.0001 | *** p : 0.001 - 0.0001 |

**e-supplementary figure 9: The format is similar to Figure 1**

#### e-supplementary Figure 10 - Arterial Blood Gas Measures

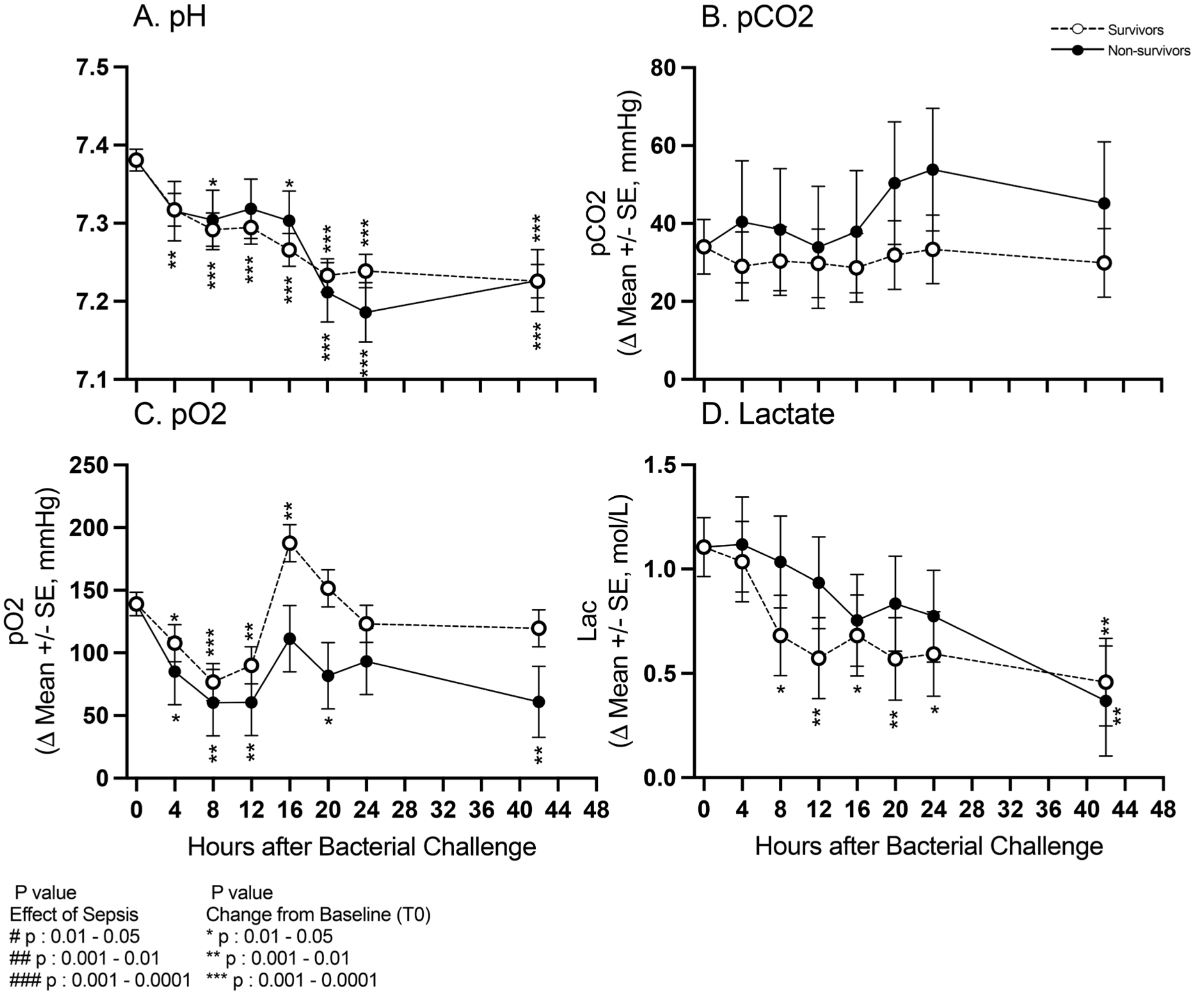


**e-supplementary figure 10: The format is similar to Figure 2**

### e-supplementary Figure 11 - Serum Chemistries

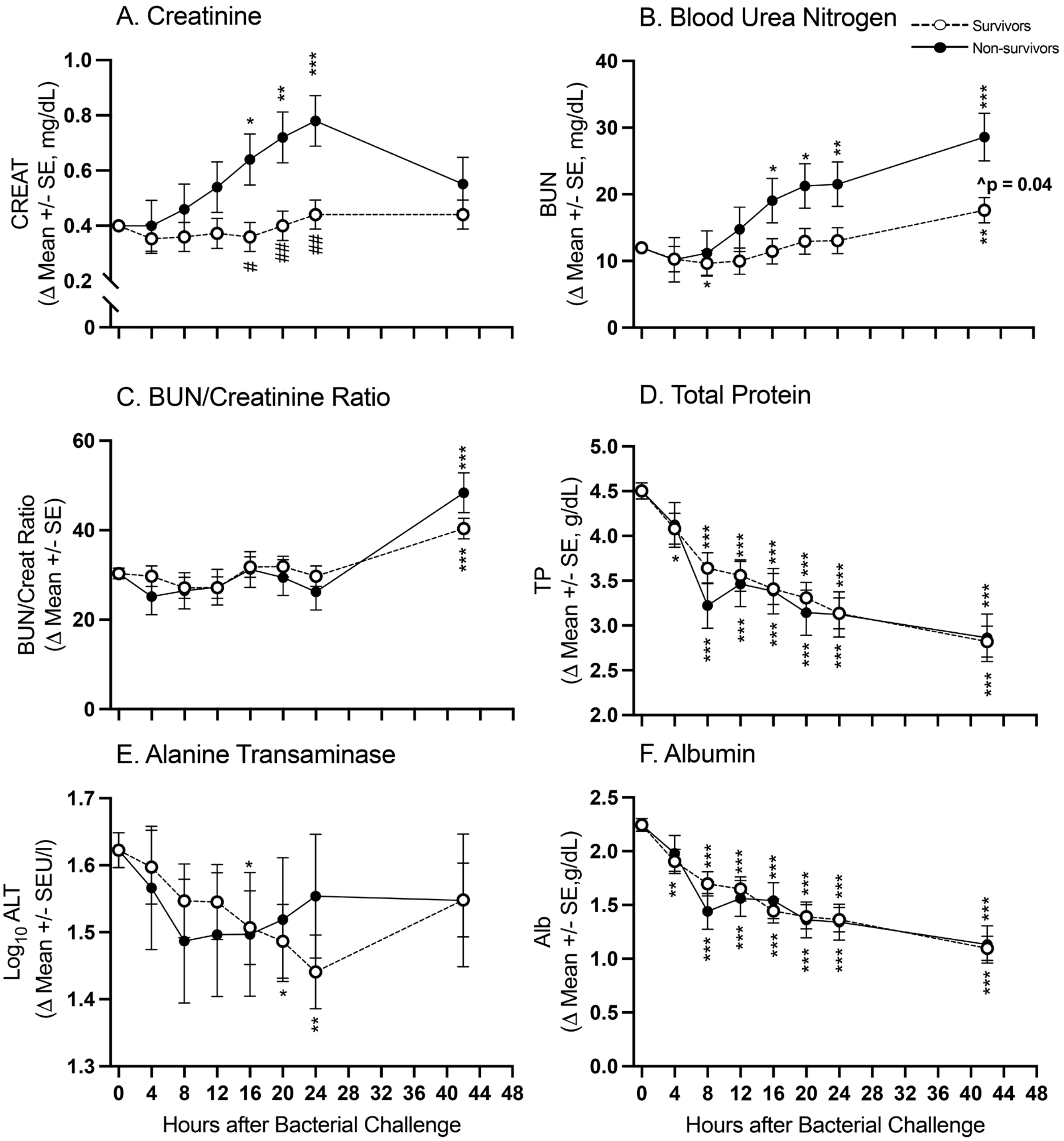


| P value<br>Effect of Sepsis | P value<br>Change from Baseline (T0) |
| --- | --- |
| # p : 0.01 - 0.05 | * p : 0.01 - 0.05 |
| ## p : 0.001 - 0.01 | ** p : 0.001 - 0.01 |
| ### p : 0.001 - 0.0001 | *** p : 0.001 - 0.0001 |

**e-supplementary figure 11: The format is similar to Figure 2**

**e-supplementary Figure 12 - Electrolytes**

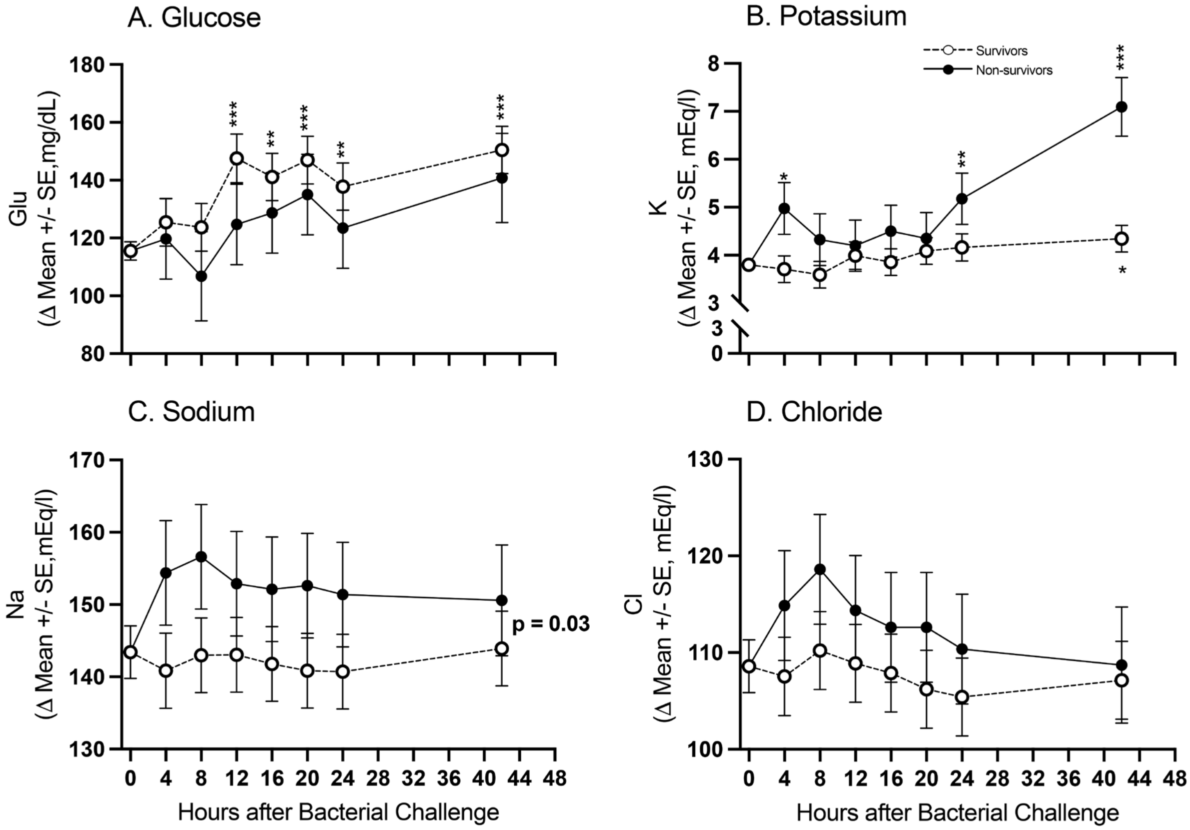


P value
Effect of Sepsis
# p : 0.01 - 0.05
## p : 0.001 - 0.01
### p : 0.001 - 0.0001

P value
Change from Baseline (T0)
* p : 0.01 - 0.05
** p : 0.001 - 0.01
*** p : 0.001 - 0.0001

**e-supplementary figure 12: The format is similar to Figure 2**

### e-supplementary Figure 13 - Complete Blood Count

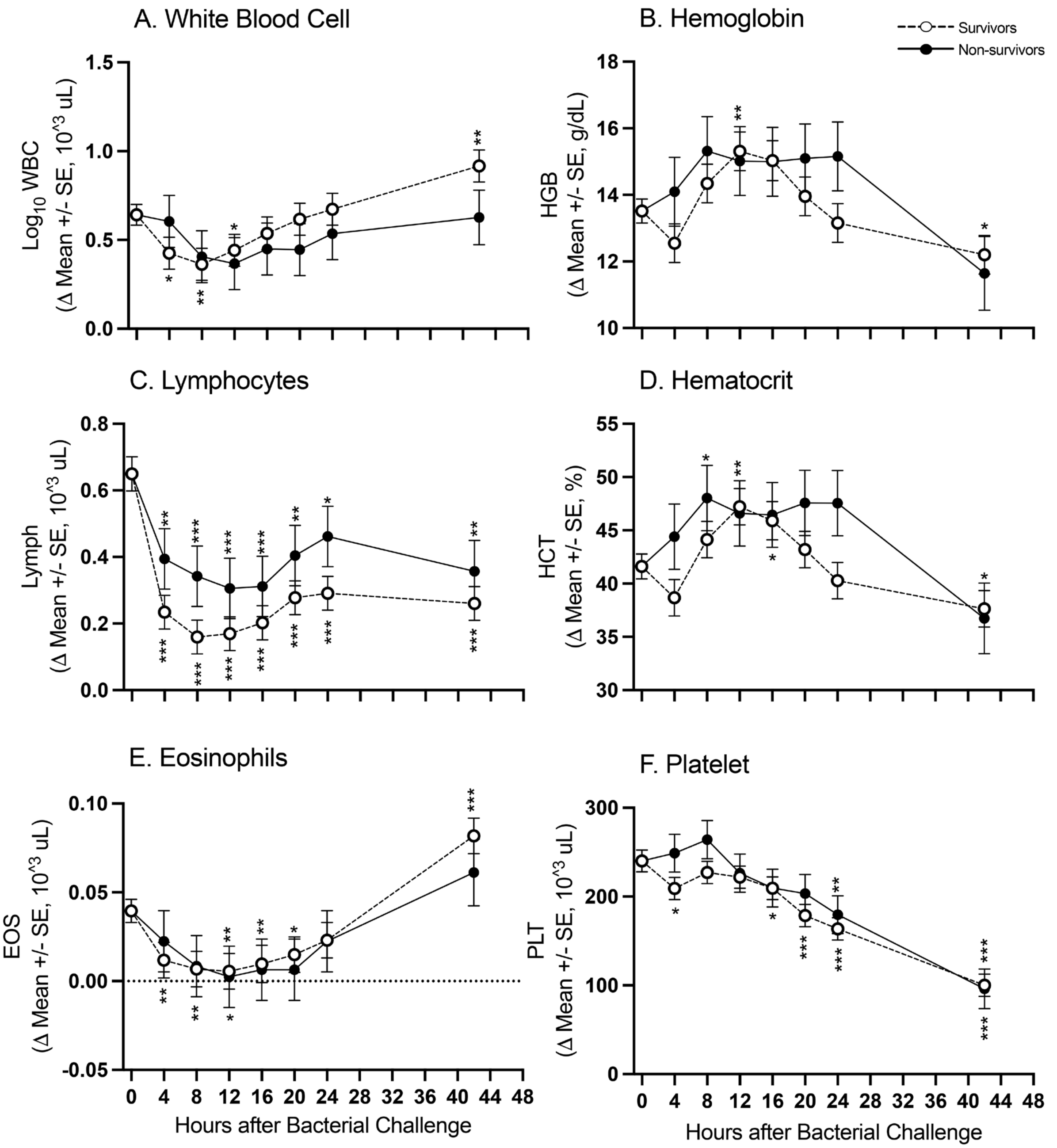


| P value | P value |
|---|---|
| Effect of Sepsis | Change from Baseline (T0) |
| # p : 0.01 - 0.05 | * p : 0.01 - 0.05 |
| ## p : 0.001 - 0.01 | ** p : 0.001 - 0.01 |
| ### p : 0.001 - 0.0001 | *** p : 0.001 - 0.0001 |

**e-supplementary figure 13: The format is similar to Figure 2**

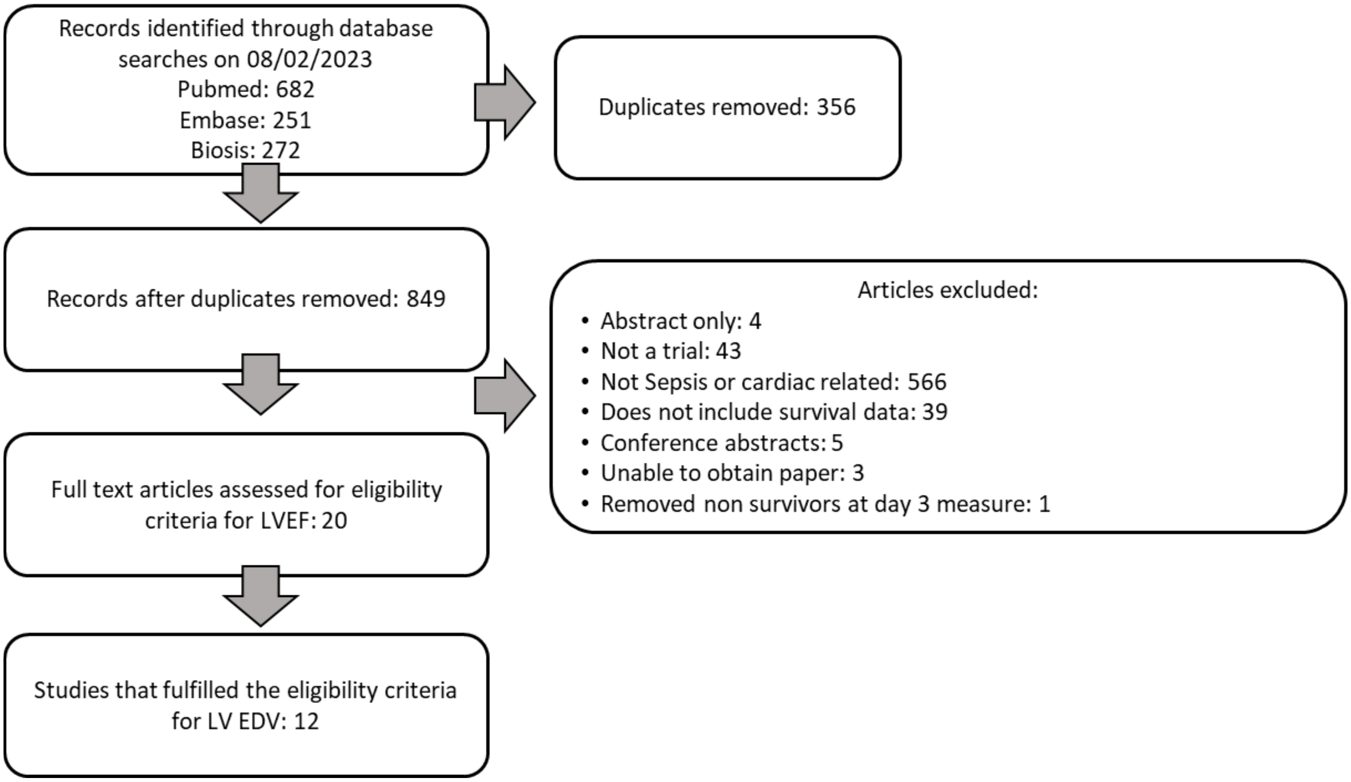


**e-supplementary Figure 14: Flow chart outlining literature review with exclusion and inclusion criteria.**

**e-supplementary table 1. Survival Summary of Treatment groups**

| Treatment Group (Total Number Studied) | **Septic (n = 27)** | **Control (n = 6)** |
|---|---|---|
| # Deaths (Time in hours from challenge) | N = 5 (29h, 65.5h, 58h, 70.5h, 73h) | N = 0 |
| # Survivors | N = 22 | N = 6 |

**e-supplementary table 2: Left Ventricular Ejection Fraction in Human Sepsis and its association with survival.**

| Study | Scan | Septic Survivor Total | Septic Survivor Mean | Septic Survivor SD | Septic Non-Survivor Total | Septic Non-Survivor Mean | Septic Non-Survivor SD | Mean Difference | Weight |
|---|---|---|---|---|---|---|---|---|---|
| 1. Spathoulas 2022 | Echo | 10 | 63.5 | 9.7 | 4 | 68.8 | 10.6 | | 1.3% |
| 2. Ehrman 2022 | Echo | 56 | 50.0 | 11.1 | 17 | 45.0 | 18.5 | | 2.1% |
| 3. Wang 2021 | Echo | 110 | 63.5 | 4.2 | 53 | 61.1 | 6.9 | | 16.3% |
| 4. Somboonviboon 2021 | Echo | 13 | 57.1 | 15.3 | 7 | 52.1 | 28.8 | | 0.4% |
| 5. Song 2020 | Echo | 37 | 34.9 | 8.7 | 12 | 32.6 | 7.5 | | 5.8% |
| 6. Lanspa 2020 | Echo | 300 | 59.9 | 13.3 | 93 | 60.6 | 14.8 | | 10.2% |
| 7. Kim 2020 | Echo | 106 | 56.5 | 13.0 | 102 | 52.3 | 14.8 | | 8.8% |
| 8. Havaldar 2018 | Echo | 29 | 30.9 | 14.3 | 19 | 36.3 | 13.9 | | 2.6% |
| 9. EL-Zayat 2018 | Echo | 21 | 52.4 | 13.2 | 29 | 51.7 | 14.4 | | 2.9% |
| 10. Zaky 2016 | Echo | 33 | 52.4 | 11.1 | 20 | 48.8 | 13.5 | | 3.4% |
| 11. De Geer 2015 | Echo | 35 | 50.0 | 11.1 | 15 | 53.0 | 16.3 | | 2.2% |
| 12. Landesberg 2014 | Echo | 65 | 56.2 | 7.8 | 41 | 56.9 | 12.3 | | 7.6% |
| 13. Harmankaya 2013 | Echo | 28 | 58.7 | 5.2 | 27 | 60.6 | 4.1 | | 14.0% |
| 14. Landesberg 2012 | Echo | 167 | 60.0 | 10.0 | 95 | 57.0 | 13.0 | | 11.5% |
| 15. Furian 2012 | Echo | 30 | 57.0 | 15.0 | 15 | 57.0 | 10.0 | | 3.1% |
| 16. McLean 2007 | Echo | 31 | 47.5 | 15.3 | 9 | 46.7 | 14.1 | | 1.6% |
| 17. Jardin 1999 | Echo | 34 | 43.9 | 16.4 | 56 | 41.6 | 10.6 | | 4.2% |
| 18. Parker 1989 | MUGA | 33 | 40.0 | 17.2 | 21 | 47.0 | 183.3 | | 0.0% |
| 19. Ellrodt 1985 | MUGA | 18 | 45.0 | 11.0 | 17 | 43.0 | 19.0 | | 1.7% |
| 20. Hoffman 1983 | MUGA | 7 | 59.0 | 17.0 | 9 | 49.0 | 24.0 | | 0.5% |
| **Summary** (Random effect model) | | **1163** | | | **661** | | | MD 1.0217 [−0.3739; 2.4174] p = 0.40, $I^2$ = 5% | **100.0%** |

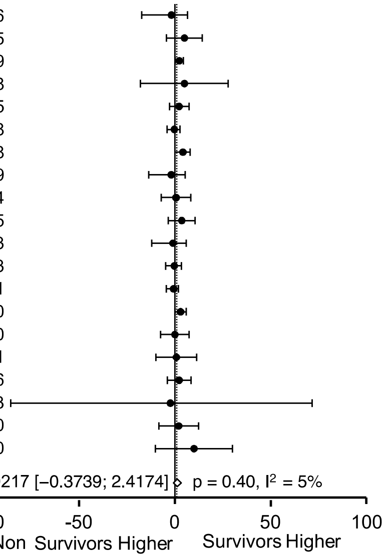


MUGA - Multigated Acquisition Scan (Radionucleotide scan)

**e-supplementary table 3: Sequential One-out sensitivity analysis for Table 1**

#### Sequential One-out Sensitivity Analysis

*(Impact of Removing Each Study on Summary p-value and $I^2$ For The Remaining 11 Studies)*

| Omitted Study | Year | Standard Mean Difference | *95% CI* | Random Effects Model p- value | Random Effects Model $I^2$ |
|---|---|---|---|---|---|
| 1. Song | 2020 | 0.1939 | [-0.3474; 0.7352] | 0.48 | 87.7% |
| 2. Havaldar | 2018 | 0.2374 | [-0.3060; 0.7809] | 0.39 | 88.0% |
| 3. Zaky | 2016 | 0.2240 | [-0.3219; 0.7698] | 0.42 | 87.6% |
| 4. Landesberg | 2014 | 0.1912 | [-0.3531; 0.7373] | 0.49 | 87.8% |
| 5. Harmankaya | 2013 | 0.2160 | [-0.3307; 0.7627] | 0.44 | 88.0% |
| 6. Landesberg | 2012 | 0.1957 | [-0.3531; 0.5162] | 0.49 | 87.5% |
| 7. Furian | 2012 | 0.1707 | [-0.3643; 0.7056] | 0.53 | 87.7% |
| 8. McLean | 2007 | 0.2147 | [-0.3275; 0.7570] | 0.43 | 88.0% |
| 9. Liu | 2000 | 0.0583 | [-0.3542; 0.4709] | 0.78 | 85.0% |
| **10. Jardin** | **1999** | **0.3638** | **[0.2115; 0.5162]** | **<0.0001** | **57.0%** |
| 11. Parker | 1989 | 0.1851 | [-0.3566; 0.7268] | 0.50 | 87.8% |
| 12. Hoffman | 1983 | 0.2817 | [-0.2301; 0.7934] | 0.28 | 87.6% |

**e-supplementary table 4: Ventricular Chamber Size in Human Sepsis is Associated with Survival.**

| | Study | Scan | Septic Survivor Total | Septic Survivor Mean | Septic Survivor SD | Septic Non-Survivor Total | Septic Non-Survivor Mean | Septic Non-Survivor SD | 95 % CI |
|---|---|---|---|---|---|---|---|---|---|
| 1 | Havaldar 2018 (ml) | Echo | 29 | 69.0 | 34.3 | 19 | 70.7 | 44.1 | [-0.6208; 0.5363] |
| 2 | Furian 2012 (ml) | Echo | 30 | 107.0 | 32.0 | 15 | 87.0 | 22.0 | [0.0380; 1.3113] |
| 3 | Song 2020 (mm) | Echo | 35 | 50.2 | 5.1 | 11 | 47.6 | 8.1 | [-0.2518; 1.1156] |
| 4 | Harmankaya 2013 (mm) | Echo | 28 | 45.0 | 5.0 | 27 | 44.0 | 4.0 | [-0.3396; 0.7206] |
| 5 | McLean 2007 (mm) | Echo | 31 | 47.0 | 10.0 | 9 | 45.0 | 9.0 | [-0.5434; -0.9436] |
| 6 | Zaky 2016 (ml/m$^2$) | Echo | 33 | 104.8 | 46.7 | 20 | 99.7 | 49.9 | [0.4509; 0.6607] |
| 7 | Landesberg 2014 (ml/m$^2$) | Echo | 65 | 56.9 | 13.4 | 41 | 51.1 | 12.7 | [0.0429; 0.8338] |
| 8 | Landesberg 2012 (cm$^3$/m$^2$) | Echo | 167 | 58.6 | 17.3 | 95 | 51.7 | 18.4 | [0.1345; 0.6427] |
| 9 | Liu 2000 (ml/m$^2$) | Echo | 15 | 72.7 | 14.6 | 10 | 42.2 | 6.6 | [1.3500; 3.5185] |
| 10 | Jardin 1999 (ml/m$^2$) | Echo | 34 | 75.3 | 20.1 | 56 | 115.0 | 25.0 | [-2.1857; -1.1953] |
| 11 | Parker 1989 (ml/m$^2$) | MUGA | 33 | 122.0 | 45.9 | 21 | 99.0 | 41.2 | [0.0434; 1.0690] |
| 12 | Hoffman 1983 (ml/m$^2$) | MUGA | 7 | 69.1 | 20.3 | 9 | 98.8 | 46.1 | [-1.7844; 0.2789] |

**Summary**

| (Random effect model) | # Studies | # Survivors | # Non survivors | |
|---|---|---|---|---|
| All Studies | 12 | 507 | 333 | $I^2$ = 87%, No summary provided |
| Outlier Removed (#10*) | 11 | 473 | 277 | SMD 0.36 [ 0.21; 0.52] $p$ < 0.0001, $I^2$ = 57% |

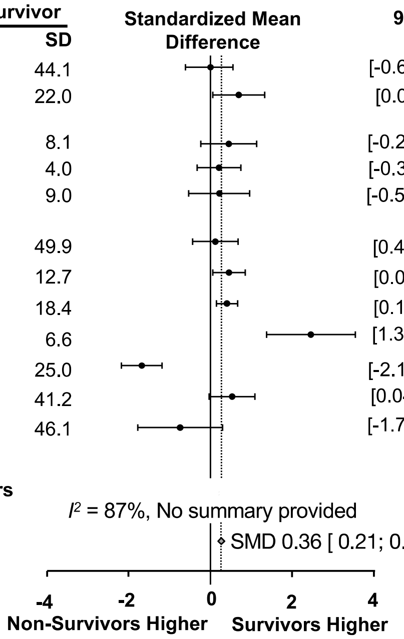


*Please see e-supplementary Summary Table 1
MUGA - Multigated Acquisition Scan (Radionucleotide scan)
